# Development of cognition in corvids

**DOI:** 10.64898/2026.02.27.708529

**Authors:** Rachael Miller, Emma Claisse, Adam Timulak, Nicola S. Clayton

## Abstract

Corvids - members of the crow family - exhibit some of the most sophisticated cognitive abilities outside the primate lineage, yet the developmental origins of many of these abilities remain poorly understood. Here, we present a systematic review of the past 20 years (from 2005) of empirical research on corvid cognitive development, synthesising evidence across core/ foundational, social and physical cognitive domains. Using a structured search strategy and detailed coding framework, we identified 47 relevant studies spanning 16 corvid species. We evaluate and discuss developmental trajectories, species/ taxa-level variation and methodological robustness across studies. For within and between-taxa comparisons, we particularly focus on the best represented abilities in the coded sample: 1) object permanence and caching; 2) tool-use/ manufacture; 3) object manipulation and play; and 4) gaze following. Corvid developmental patterns show both parallels and divergences from those documented in primates and other taxa. However, the existing corvid evidence base is constrained by small samples, captive biases, limited longitudinal data and under-representation of key cognitive abilities, such as executive function, causal reasoning, self-control, metacognition, spatial memory and social learning. We outline critical gaps and future directions, emphasising the need for comparative, longitudinal and ecologically grounded approaches, including the science of magic and Theory of Mind, to better understand how early-life cognition shapes later behaviour, cognition and fitness in this model avian family.

## 1. Introduction

Cognition is broadly defined as the mental processes underlying perception, learning, decision-making and memory, which drives behaviour, and the ability to adapt to novel, human-imposed environmental challenges (Shettleworth, 2010). Since the early years of the field’s conception, primates have been at the forefront of animal cognition research. As these early studies aimed to uncover the evolutionary roots of human intelligence, primates (in particular the great apes), were subject to new research interest due to their close genetic relationship to humans.

Discoveries by pioneering scientists showed that many abilities once thought to be unique to humans, such as complex problem solving (Kohler, 2018), tool use and manufacture (Goodall, 1986) and complex social understanding (Premack & Woodruff, 1978), were in fact shared with our primate cousins, suggesting that these abilities may have deeper evolutionary roots than previously thought. However, whilst influential, this anthropocentric approach neglected to capture the full spectrum of animal intelligence and therefore its evolution across taxa.

A major step forward in this respect occurred in the late 20th century, when animal cognition researchers widened their scope and discovered the fascinating mental capabilities of an underappreciated group of birds, the corvids (members of the crow family). Although admired for their intelligence throughout the world in folklore, such as the three-legged crow of Japan and China, corvids (and birds in general) were not previously given sufficient research attention and were deemed to be unintelligent in popular culture (i.e., ‘bird brain’). However, over the next few decades, corvids quickly demonstrated their advanced cognitive complexity, displaying many abilities that were previously only apparent in the great apes (Emery & Clayton, 2004). As well as showing a particularly high degree of brain encephalisation (ratio of brain size to body size), comparable to that of great apes (Olkowicz et al, 2016), corvids produce a higher frequency of complex behaviours, such as foraging innovations and social practices, than any other bird group (with parrots close behind) (Taylor, 2014). Corvid research findings revealed that complex cognition is not restricted to the primate lineage and has indeed arisen convergently within birds. Accordingly, this research has the potential to shed light onto the mechanisms underpinning the evolution of intelligent minds, as we discover what factors in an animal’s life (including our own) lead to selection for advanced cognition.

The cognitive abilities of the various corvid species stretch across core, physical and social domains, and even transcend subjective time (Emery & Clayton, 2004). Whilst Clark’s nutcrackers, *Nucifraga columbiana*, are able to remember around 10,000 locations where they had previously stored food, due to an exceptional long-term spatial memory ability (Balda & Kamil, 1992), New Caledonian crows, *Corvus moneduloides*, are one of the best tool users and manufacturers in the non-human animal kingdom (Hunt, 1996). Even corvid species that do not use tools in the wild, such as rooks, *Corvus frugilegus*, show tool use and modification (Bird & Emery, 2009a) and appear to understand the underlying causal mechanisms of these tools (Seed et al, 2006). Often living in complex social systems, corvids also excel in social intelligence (Clayton & Emery, 2007). Ravens, *Corvus corax*, appear to understand that other individuals have their own ideas, beliefs and desires (theory of mind) (Bugnyar et al, 2016). Jackdaws, *Coloeus monedula*, recognise individuals of other species (humans) (Davidson et al, 2015) and learn from others who to avoid (Lee et al, 2019). Pinyon jays, *Gymnorhinus cyanocephalus*, are able to draw sophisticated inferences about their own dominance status based on interactions between other known and unknown individuals (transitive inference) (Paz-y-Miño et al, 2004). Furthermore, corvids, including Eurasian jays, *Garrulus glandarius*, and scrub jays, *Aphelocoma californica*, have been shown to travel through their own subjective time (mental time travel) to recall the past (Davies et al, 2024; Clayton & Dickinson, 1998) and to predict the future (Boeckle et al, 2020; Cheke & Clayton, 2012), demonstrating (before even great apes) that many animals are not ‘stuck in time’ as once thought.

However, despite the fact that decades of study have revealed that some of the most common and widespread corvid species (i.e., carrion crows, Eurasian jays, jackdaws, rooks and common ravens) possess these “complex” cognitive abilities (Bird & Emery, 2009b; Bugnyar et al, 2016; Cheke & Clayton 2012; Davidson et al, 2015; Davies et al, 2024; Emery & Clayton, 2004; Seed et al, 2006; Smirnova et al, 2015), research focusing on the ontogeny or development of cognition in corvids remains relatively rare. This represents an important knowledge gap, as “core” or “foundational” cognition underlies more complex abilities, and development/ ontogeny is a fundamental proximate explanation for understanding animal behaviour (Manning & Dawkins, 2012). For example, object permanence ability (recognising that objects continue to exist even though they are no longer visible; Ramsay & Campos, 1978) lays the foundation for memory and caching (hiding food and other items for later recovery) behaviour. While the development of some abilities, like object permanence, have been studied in several corvid species, indicating evidence of “Stage 6” level (invisible displacement) understanding (Zucca et al, 2007), many other abilities require future focus. This is likely due to the logistical, funding and ethical considerations of testing corvids from juvenile to adulthood, including the costs of housing these long-lived birds. This knowledge is crucial for informing and addressing related questions, such as how early-life cognition supports social living, cooperation, communication and cultural transmission, and underpins later complex behaviours, like perspective-taking, caching strategies and recognition of others. Furthermore, this underlies assessment of how cognitive development may relate to and impact on animal welfare or conservation outcomes, including fitness, like reproduction and survival.

In this article, we systematically reviewed the development of cognition in corvids across the past 20 years (from 2005), with the aims of broadly identifying: 1) what we currently know about developmental patterns, processes and influences on corvid cognition, including cognitive performance in juveniles and sub-adults; and 2) what we are missing in the current literature and the focus of future research, including under-represented cognitive domains or abilities. We focus on cognitive domains relating to core/ foundational, social, physical/ technical cognition, and assess specific related abilities and paradigms within these domains. We outline the four key objectives in Table 1.

### Objectives

**Table 1.**
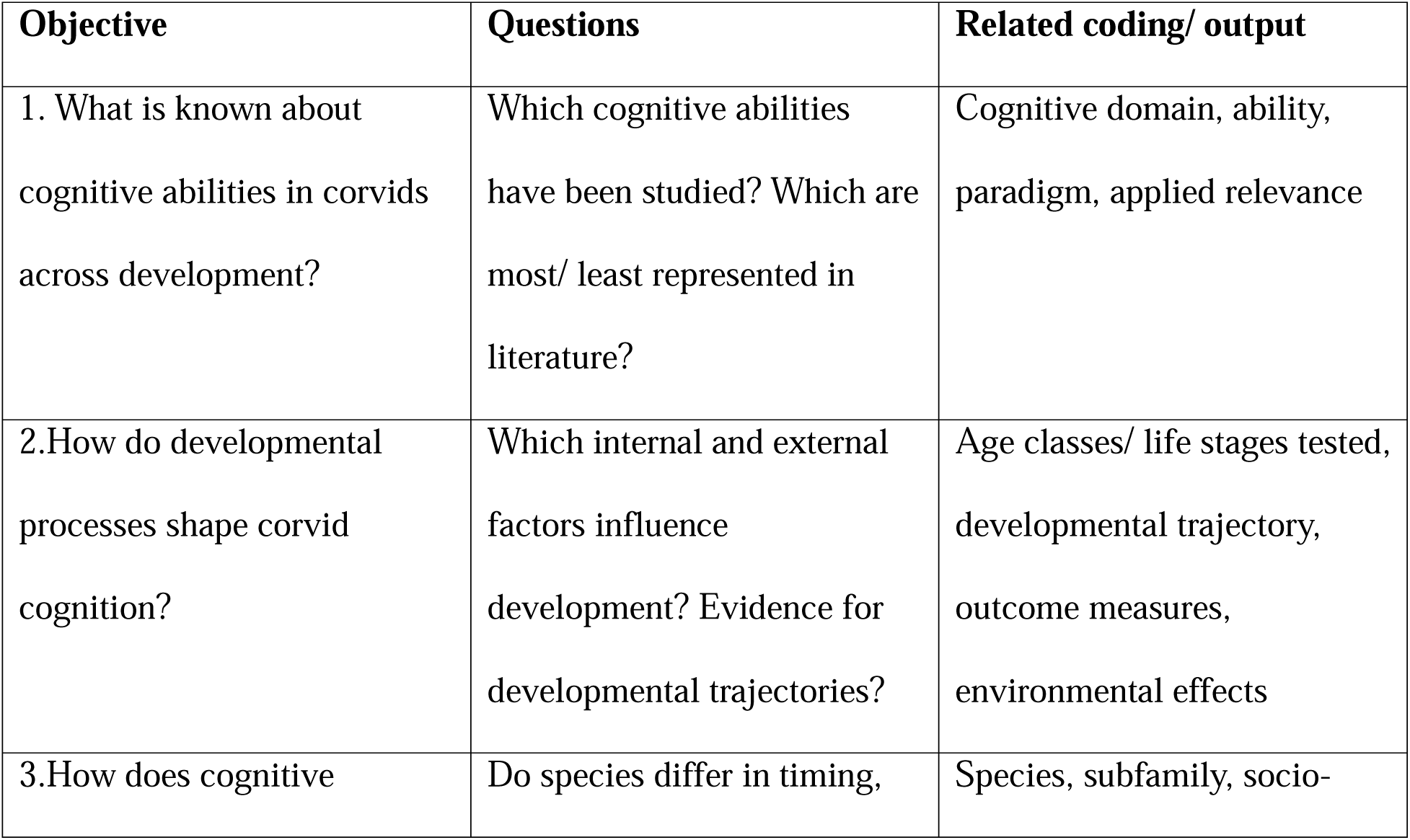
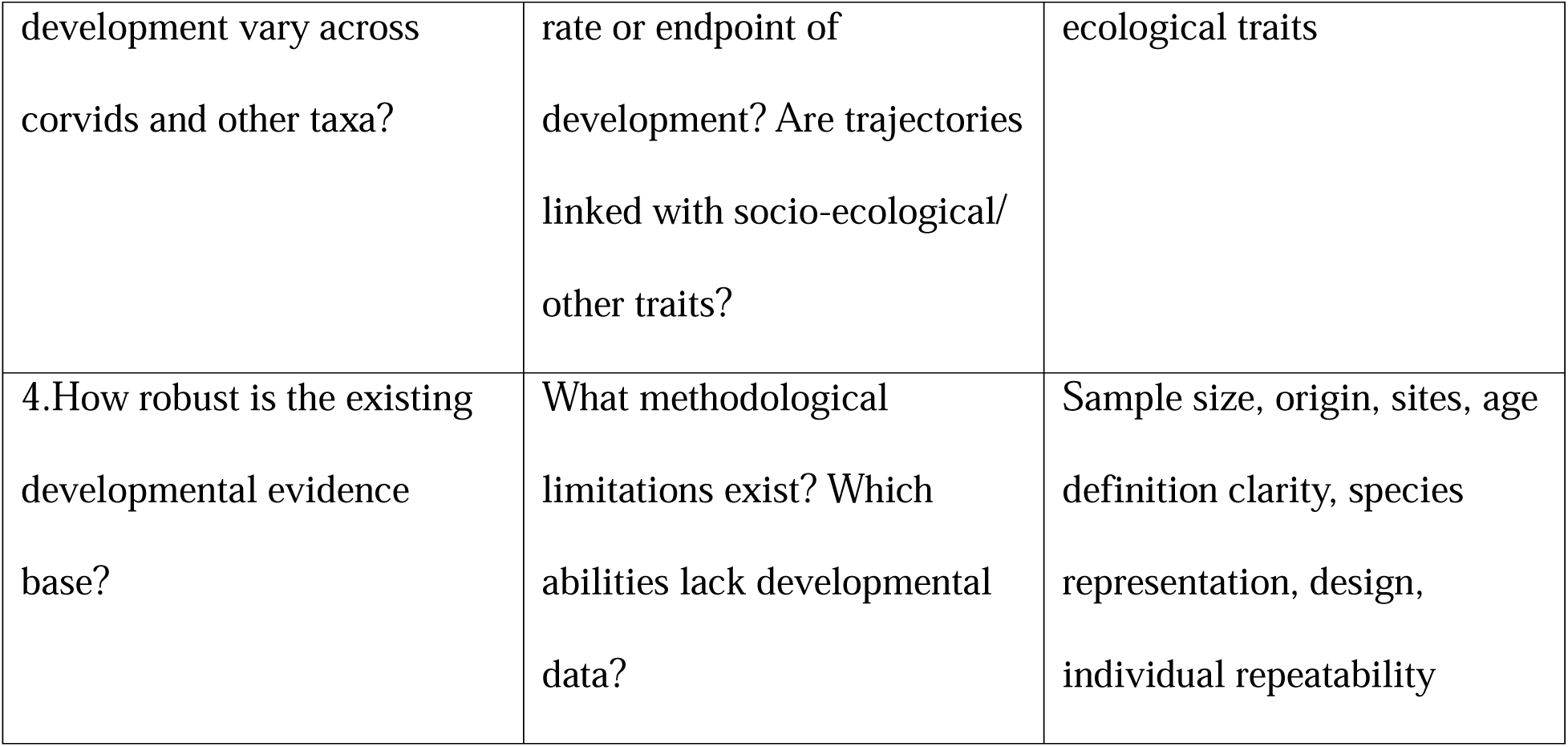
Objectives, research questions, and related coding/ outputs.

## 2. Review Methods

### Article selection process

We used a systematic review and conducted two keyword searches, outlined in detail below, with the aim of capturing a broad overview of the relevant literature relating to “corvid cognitive development” in the past 20 years.

**Keyword search 1:** ((TS=(corvid* OR corvus OR crow* OR raven* OR jay* OR rook* OR magpie* OR jackdaw* OR chough* OR treepie* OR nutcracker*) AND TS=(cogniti* OR intell* OR psycholog*) AND TS=(physical OR social OR technical OR mental OR core OR foundation*) AND TS=(cogniti* OR intell* OR memory OR learn* OR atten*) AND TS=(devel* OR ontogen*)))

**Search 1 (corvid, cognition, development):** Using keyword search 1 above (including learn, memory, atten) in the Web of Science (WoS; Clarivate) database on 12 December 2025, resulted in 1,518 articles, filtering by “research article” reduces this to 1,122 articles. In WoS, we then filtered by research area, with the aim of focusing on cognitive/ behavioural research only, to select the following: psychology (217), neurosciences neurology (75), behavioral sciences (50), zoology (48), evolutionary biology (10), veterinary sciences (5), acoustics (1), biodiversity conservation (2), developmental biology (1), reducing the articles to 318. We further filtered by publication year to include 2005-2025, as the majority of articles were from this period (original dates covered papers from 1978) and were deemed more likely to include empirical data available for extraction/ comparison. This reduced the output to 272 articles.

**Keyword search 2:** ((TS=(corvid* OR corvus OR crow* OR raven* OR jay* OR rook* OR magpie* OR jackdaw* OR chough* OR treepie* OR nutcracker*) AND TS=(developm* OR ontogen*)))

**Search 2 (corvid, development, not cognition):** Using keyword search 2 above (not including cognition or related terms) in the WoS database on 8 December 2025, resulted in 34,984 articles, filtering by “research article” reduced this to 28,251 articles. In WoS, we then filtered by research area, with the aim of focussing on cognitive/ behavioural research only, to select the following: psychology (796) and zoology (1031), further reducing the articles to 1,781. We further filtered by publication year to include 2005-2025, as the majority of articles were from this period (original dates covered papers from 1901). This reduced the output to 1,252 articles.

### Combining search outputs

For the output of search 1 and 2, using excel spreadsheets, we manually searched to include only articles that focused on: a) corvid(s) or specific corvid species (in title or abstract) as subjects, b) cognition or behaviour (including learning, memory, attention; in title or abstract), c) ontogeny, development, age effects or testing in sub-adults or juvenile birds (in title, abstract and paper), d) empirical data (i.e. not a review/ commentary/ perspective). This reduced the output to 20 articles for search 1 and 38 articles for search 2, resulting in 39 unique articles when combining both searches.

### Cross-checking against Google Scholar searches

Finally, to ensure there was good representation and coverage of the relevant empirical literature in our final sample, we cross-checked our final WoS output on a selection of manually searched articles identified from Google Scholar. We used the search terms “cognitive development corvids” and “ontogeny of cognition in corvids” and manually searched for papers covering “development/ ontogeny/ age effects” in 1) Google Scholar generally, and 2) Google Scholar profiles (with full publication lists) of fifteen established, current corvid lab leaders across the world (e.g. including ten lab leaders from Miller et al, 2022). The vast majority of articles were represented in our final WoS keyword search output, thus validating our WoS keyword search terms and supporting efforts to capture a broad representation of relevant empirical literature from 2005-2025. We identified eight further papers in this final cross-checking step (for example, published papers in journals that are not indexed, like *Animal Cognition & Behavior*), resulting in a total of 47 articles for coding.

### Summary of coding for systematic review (47 articles)

We (RM, EC, AT) first coded 10% of the same articles (5 articles) for inter-observer reliability checking, before we proceeded to code the remaining 42 articles. We coded the final articles based on the following eight categories, relating to our four key objectives (outlined in Table 1): a) study metadata (e.g. journal, years, setting), b) species and life history (e.g. species, origin, life history), c) sample and developmental staging (age, sex), d) context and experience (e.g. housing), e) cognitive domain, ability and task (e.g. paradigm, outcome measure), f) developmental patterns and mechanisms (e.g. developmental pattern, repeatability), g) Tinbergen’s questions (core components of ontogeny, mechanism, function, phylogeny) and h) quality and relevance indicators (e.g. sample size risk, age definition clarity, conservation/ welfare relevance). The full coding definitions are outlined in S1 Text.

## 3. Review Results

### Objective 1: What is known about cognitive abilities across development in corvids?

Of the 47 papers coded in our systematic review, approximately 32% (12 papers as core; 3 papers as core/ physical or core/ social) were coded as primarily core/foundational, 13% (6) were physical, 34% (16) were social and 21% (10) were social/ physical cognition domain focused.

These papers broadly covered topics/ abilities including object permanence, caching, associative learning, spatial memory, motor self-regulation, gaze following, neophobia/ exploration, object manipulation, tool-use, problem-solving, causal reasoning, communication, social behaviour, social learning and play. Commonly used paradigms or methods included: novel/ familiar stimuli/ object manipulation or combinations, behavioural observations, video/ image presentations, acoustic recording, acoustic playbacks, caching/ foraging, radial arm maze, detour, tool-use/ manufacture, trap-tube and associative/reversal learning tasks. S1 and S2 Tables provide a summary of the most commonly tested domains and abilities across the 16 represented corvid species. Of the 16 tested species, 69% (11) of species were tested on core/ foundational, 56% (9) on social and 31% (5) on physical cognition domains.

Only 3 papers in our sample had clear applied relevance for conservation or welfare, with a further 10 papers coded as “implicitly” (i.e. not directly) indicating more applied relevance. 40 papers included a developmental focus (for 5 of which, this was only “suggested”), 6 papers included a focus on neuro/hormone/physiology, 7 papers included fitness influences (for 6 of these, this was only “suggested”) and 7 papers tested multi-species. The study countries where data collection took place were: 21 papers in Austria, 8 in USA, 5 in UK, 4 in New Caledonia, 3 in Canada, 3 in Sweden, 3 in Germany, 2 in China, 1 in Italy, 1 in Russia and 1 in Spain.

### Objective 2: How do developmental processes shape corvid cognition?

Within our coded output, the reported age classes used were: 72% (34) papers with juveniles, 17% (8) nestlings, 32% (15) fledglings, 26% (12) sub-adults and 30% (14) adults, with most papers testing subjects across age classes. For the 33 papers that identified developmental trajectories, 61% (20) papers reported “improves with age”, 12% (4) as “declines with age”, with 15% (5) as “stable” and 12% (4) as “no pattern” (14 papers did not analyse developmental patterns). The summary of related coded output can be found in S3 Table, with the full coding output available via Figshare: 10.6084/m9.figshare.31410981 (private link: https://figshare.com/s/b88ae401b55b99b35d57). Social behaviour, such as interactions, social structure, antipredator behaviour, social foraging, social learning and communication were the most common types of social “cognition” tested (S3 Table). Across social cognition related papers, there did not appear to be a consistent developmental trajectory across species found for general social behaviour measures, at least in terms of how this was reported - and thus coded by us - in the individual papers.

In the following section, we present the research findings in more detail in relation to the best represented abilities in the coded sample: 1) object permanence and caching; 2) tool-use/ manufacture; 3) object manipulation and play; and 4) gaze following.

#### Object permanence

Piaget posited that during the first two years of life, also known as the sensorimotor stage, human children develop this foundational concept in a fixed sequence of six stages, which is driven primarily by internal maturation rather than experience or learning (Piaget, 1952, 1954; Table 2). Stages 1 and 2 focus on awareness and tracking of objects, while Stages 3 through 6 involve the ability to search for occluded objects with increasing complexity. Most animal cognition research is concerned with Stages 4 through 6, which involve retrieval of fully occluded objects and visible/ invisible displacements. Achieving Stage 6 object permanence is generally taken to indicate a flexible mental representation of objects that are no longer perceptually available (Ramsay & Campos, 1978).

**Table 2.**
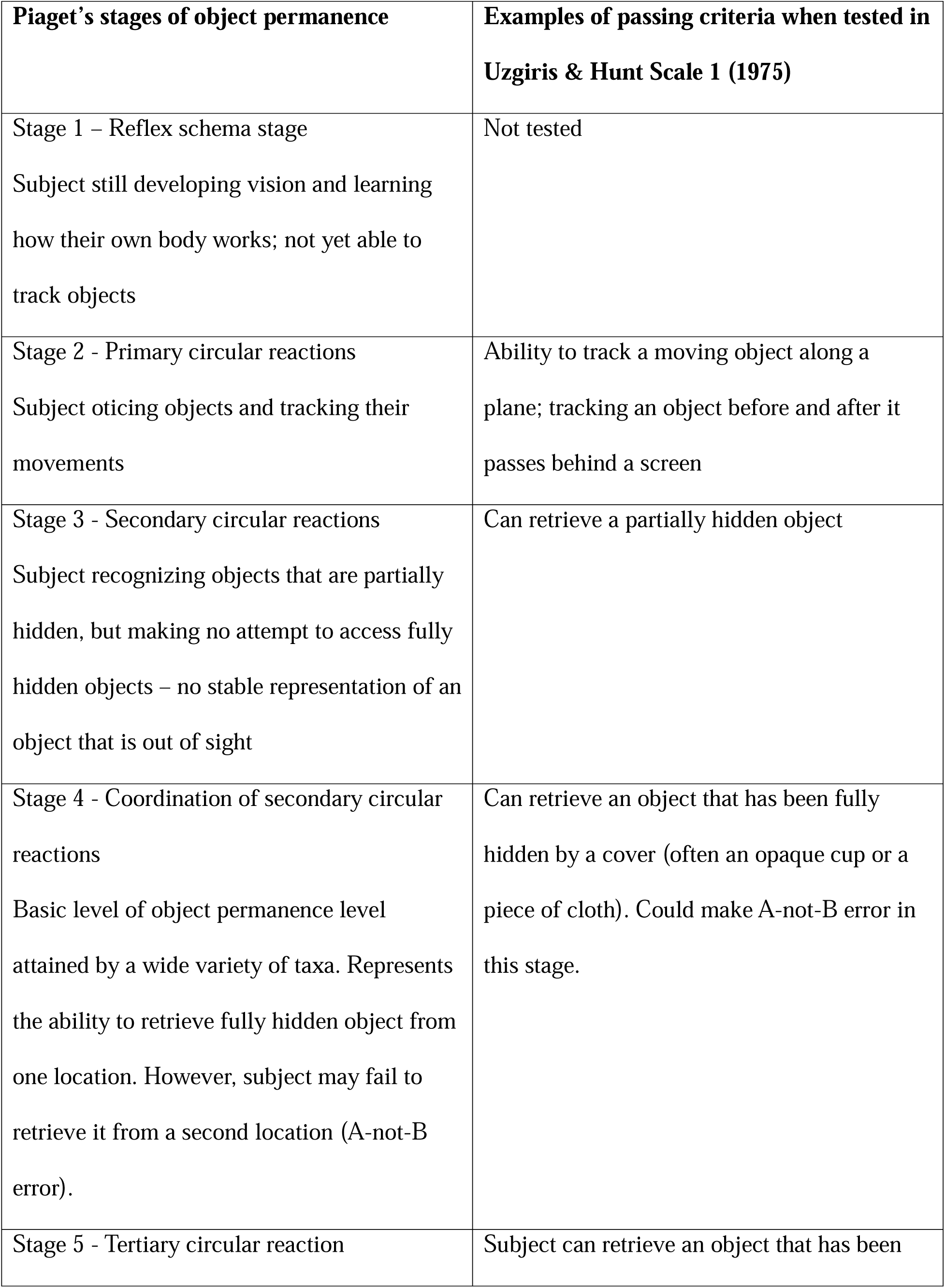
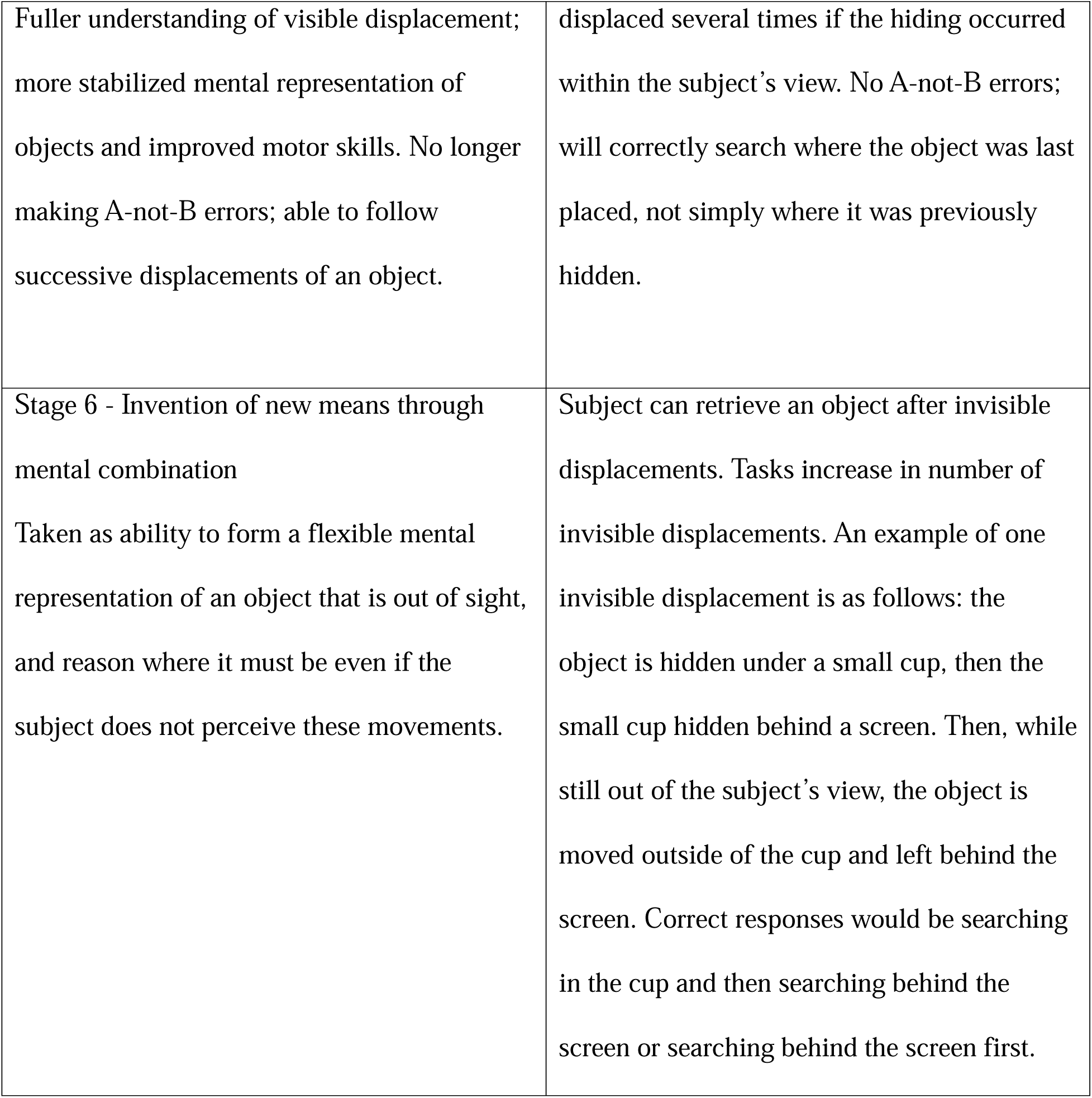
An overview of Piaget’s six stages of object permanence and the associated tasks in Uzgiris and Hunt’s Scale 1 (1975)

Since the 1970s, Uzgiris and Hunt’s Scale 1 tasks within the Piagetian framework have been widely used to assess object permanence across non-human taxa (Table 2). Using this scale, representatives of the great apes (Barth & Call, 2006; de Blois et al, 1998), parrots (Funk, 1996; Pepperberg & Funk, 1990; Pepperberg & Kozak, 1986; Pepperberg et al, 1997) and corvids (e.g. Pollok et al, 2000; Zucca et al, 2007) have demonstrated competence up to Stage 6, including invisible displacements. The scale provides a series of 15 incremental, non-verbal tasks that can be performed with simple materials, each corresponding with a Piagetian Stage - a standardized framework that has facilitated direct cross-species and cross-study comparisons. The first studies to apply Scale 1 to avian ontogeny emerged in the 1990s (Funk, 1996; Pepperberg et al, 1997).

Over the last 25 years, the development of object permanence in juvenile corvids has been examined in Eurasian magpies, *Pica pica* (Pollok et al, 2000), common ravens (Bugnyar et al, 2007), carrion crows, *Corvus corone* (Hoffmann et al, 2011), jackdaws (Ujfalussy et al, 2012), Eurasian jays (Zucca et al, 2007), California scrub-jays (Salwiczek et al, 2009) and azure-winged magpies, *Cyanopica cyanus* (Wang et al, 2021) (Fig. 1; S4 Table). Rooks have also been recently assessed for object permanence, although not from an ontogenetic perspective (i.e. in adult birds only; Cornero & Clayton, 2025).

**Fig. 1.**
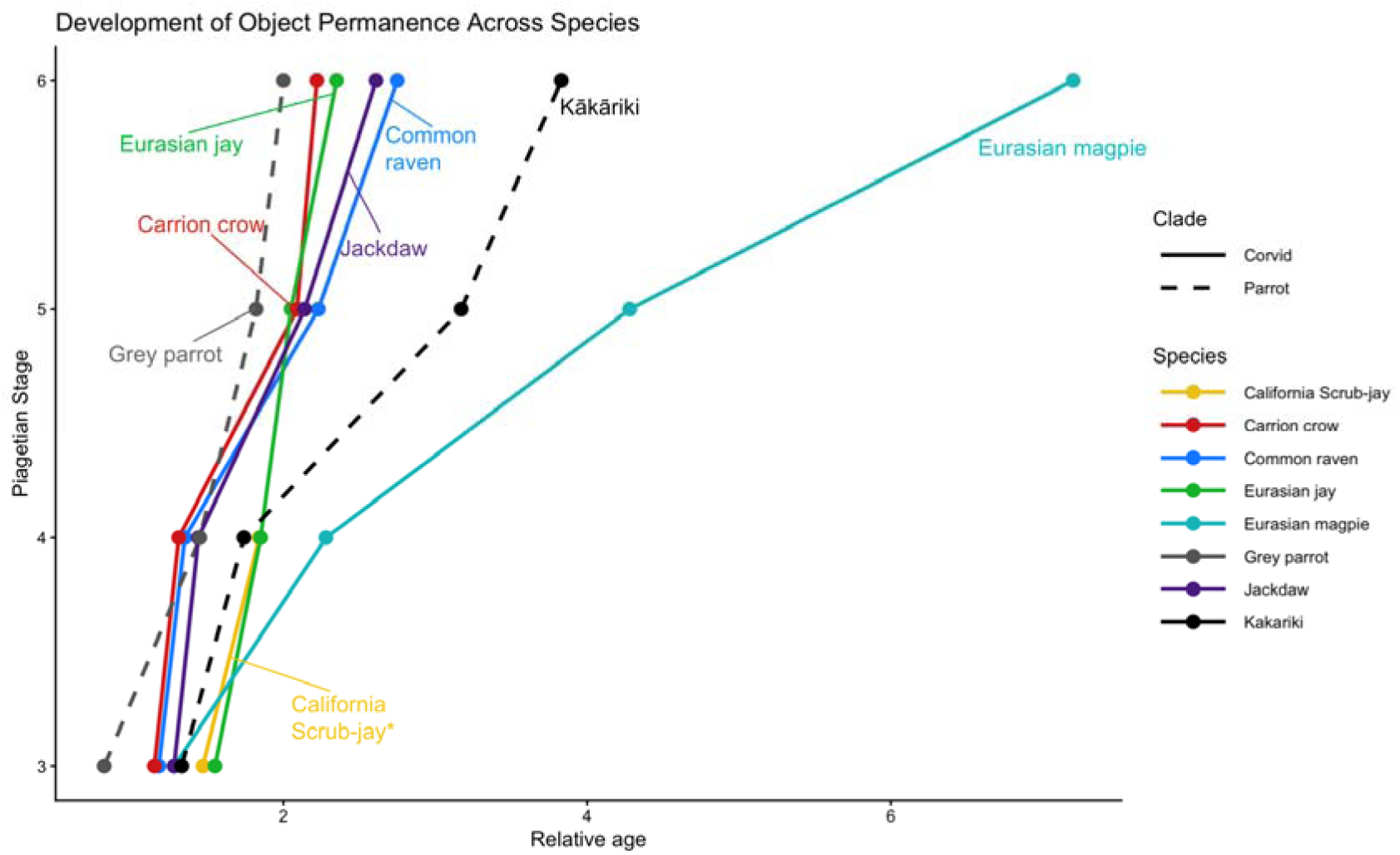
Development of object permanence across corvid and parrot species using data from Bugnyar et al, 2007; Hoffmann et al, 2012; Funk, 1996; Pepperberg et al, 1997; Pollok et al, 2000; Salwiczek et al, 2009; Ujfalussy et al, 2012; Zucca et al, 2007. S4 Table contains the data used to make this figure; note Task 15 is excluded in the figure

To make this comparison, we divided the age at development (in days), by the average age at fledging for the species. Age at fledging was the chosen unit to represent the length of the developmental period for each of the species, as Hoffmann et al (2011) did for their comparison of object permanence in corvids. When controlling for hatching to fledging time in this way, we found that grey parrots and kākārikis had a similar trajectory to the corvids, excluding magpies. As Hoffmann et al (2011) noted, magpies’ slower accomplishment of Scale 1 tasks (Pollok, 2000) stands out given their ecological similarities to the other species tested, including food-caching behaviour, overall generalist diet and variable social environments. This apparent difference in performance may be due to methodological differences like frequency of testing.

Outcome measures across ontogeny studies were typically the age at which subjects were successful at each task on the scale and achieved each associated stage, however criteria for “passing” a task varied per study. With the exception of California scrub-jays, which were only tested for Stages 1-4 (Salwiczek et al, 2009), all corvid species tested have notably shown at least some evidence for Stage 6 competence, although not all subjects in the experiments completed every task in the scale. Later tasks in the sequence were typically more difficult, and no carrion crows were able to complete Task 15 (Hoffmann et al, 2012).

Across studies, there were also pronounced interspecific and interindividual differences in the occurrence of “a-not-b” errors - continuing to search for an object in a previously correct location even after observing the object be hidden in a new location, which become relevant in the tasks following Stage 4 achievement - retrieval of a fully occluded object. In Task 5 on Uzgiris & Hunt’s Scale 1, which is the first of several tasks associated with Piagetian Stage 5 object permanence, an object is repeatedly hidden in one location several times before being visibly displaced to a new location. Success requires searching for the object where it was last placed, rather than where it was rewarded in the previous trials. Inhibitory control - to control reflexive, conditioned or otherwise learned responses in order to choose a conflicting, more rewarding or complex course of action (Miller et al, 2019) - not only the understanding of object permanence, has been suggested to influence the occurrence of these errors. While the A-not-B task has been widely used as a measure of self-control (Maclean et al, 2014), as it involves inhibiting a previously rewarded response, an intervention study on adult New Caledonian crows provided evidence to suggest poor performance may be impacted by other factors (Jelbert et al, 2016). After all 14 crows initially failed at the A-not-B test, Jelbert et al (2016) trained one group (1) to attend to the movements of food in a human hand as it baited one of three cups and covered them all with lids, and a second group (2) on a task designed to train inhibitory control without the involvement of human hands or the movement of a reward. After this intervention training, only the hand-tracking group (1) improved their performance on the A-not-B task, indicating that performance in Scale 1 tasks may reflect the level of proficiency in tracking human hands rather than inhibitory control. These findings raise broader concerns about the extent to which factors such as embodiment differences, associative learning, executive function and experimenter cueing may influence performance on Scale 1 object permanence tasks.

#### Caching

The development of object permanence is closely linked with caching - a widespread behaviour across the corvid family (de Kort & Clayton, 2005). Although caching varies considerably between species in frequency, number of caches and duration of storage, the ability to hide food out of sight and retrieve it later implies at least a rudimentary understanding of object permanence. Additionally, cache pilfering (stealing others’ caches) also relies on object permanence, as it involves tracking and remembering the hidden food of others. Furthermore, caching behaviour involves episodic-like memory, in which individuals remember the what-where-when of specific events (Clayton & Dickinson, 1998; Davies & Clayton, 2024) and can also keep track of who was watching when (Dally et al, 2006).

Two ontogenetic studies explicitly examined the development of caching behaviour alongside performance on Scale 1 object permanence tasks (Bugnyar et al, 2007; Salwiczek et al, 2009). Together, these studies indicate a temporal relationship between the maturation of object permanence and the acquisition of caching behaviour. In both common ravens and Californian scrub-jays, basic motor components of caching, such as placing or inserting objects into substrate, emerged before individuals were able to independently retrieve fully hidden items.

Functional or tentative caching, however, only appeared after or around the time individuals succeeded on Piagetian Stage 4 tasks, when they could reliably retrieve completely occluded objects. These findings suggest that while the motor components of caching may precede full object permanence, successful caching is related to the acquisition of at least Stage 4 competence, which occurred experimentally at approximately 35 days in scrub-jays (Salwiczek et al, 2009) and 54 days in ravens (Bugnyar et al, 2007). While caching implies object permanence, and the two appear developmentally linked, it is also important to note that full object permanence capabilities have been documented across a variety of non-food-storing taxa (e.g. parrots, Pepperberg & Funk, 1990; Pepperberg & Funk, 1990; great apes, see Table S5; Asian elephants, *Elephas maximus*, Miller, 2019), suggesting object permanence is a broadly adaptive cognitive trait that supports many ecological demands beyond caching.

Social cognition also plays an important role in caching, with decades of research demonstrating flexible caching techniques that take into account the presence and knowledge of conspecifics (or potential competitors) (Bugnyar & Kotrschal, 2002; Clayton et al, 2007; Emery et al, 2004; see Grodzinski & Clayton, 2010 for a review). Furthermore, it appears that caching techniques are mediated by lived experience. For example, Emery & Clayton (2001) allowed adult scrub-jays to cache either in private or while observed by a conspecific, and found the birds subsequently re-cached food at new sites during recovery trials only if they had prior experience pilfering others’ caches. The cache-protection and tactical deception behaviours that adult corvids display, specifically re-caching after being watched, have also been considered potential evidence of theory of mind (Bugnyar et al, 2007 for a review in ravens).

There has been comparatively little research on social cognition and caching in juvenile birds specifically. However, the limited developmental evidence available suggests that caching-related social behaviours continue to be shaped substantially after fledging. Bugnyar and colleagues (2007) concluded that in the months after juvenile ravens fledge, the positioning of caches relative to an observer and the use of cache protection tactics like aggression or recovery, is influenced by experience. Miller et al (2023) also found that social attention in caching context is influenced by development, with juvenile ravens and carrion/hooded crows showing higher use of barriers than fledglings or sub-adults. In field settings, the development of caching location choice and social context has been recently evaluated in ravens (Beck et al, 2020) and Florida scrub-jays, *Aphelocoma coerulescens* (Fuirst et al, 2020), also finding that experience has influence, though specific mechanisms are difficult to identify solely through observation.

While existing works indicate that social elements of caching are gradually shaped by experience, the more specific developmental pathways that lead to sophisticated and flexible adult caching behaviour remain relatively underexplored in the existing literature.

#### Tool use/ manufacture

Kenward et al (2011) proposed that caching may be an evolutionary precursor to tool use - a topic which is similarly underexplored in the existing ontogenetic corvid literature. Of the seven tool use studies in our coded sample, six focused on New Caledonian crows - the only corvid habitual tool user currently known with a wild population (Hunt et al, 2007; Rutz & St Claire, 2012 – note that Alalā are also habitual tool users but currently are Extinct in the Wild with ongoing reintroduction efforts). The other two taxa represented in the sample were rooks (Tebbich et al, 2007) and ravens, who were also compared with New Caledonian crows (Jacobs & Osvath, 2023). Both rooks and ravens are capable of using tools to solve problems in captivity (examples of non-developmental studies with sub-adult/ adult corvid: rooks, “trap-tube” task: Seed et al, 2006; “Aesop’s fable water displacement” task: Bird & Emery, 2009b, as are jays: Cheke et al, 2011), though these species’ are not known to use tools in the wild.

The developmental studies on New Caledonian crows indicate that tool use consistently improves with age (S3 Table). Holzhaider et al (2010b) suggested that there may be a sensitive period early in the life of the New Caledonian crow, where they would likely observe and may remember tool manufacture displayed by their parents. This would be followed by the juveniles practicing this behaviour until they are proficient at manufacturing the tools themselves. Indeed, Holzhaider et al (2010b) found that juveniles would often attempt to use their parents’ discarded tools from 2 months old, the earliest tested age, to 6 months old.

Ontogenetic studies on habitual tool users, like New Caledonian crows, are of particular interest because they provide insight into the role of social or individual learning versus an animal’s inherent propensity for making tools or a genetic basis. Kenward et al (2006) found that juvenile New Caledonian crows showed motor patterns with object manipulations (tool-oriented behaviour) that resembled tool use/manufacture displayed by adults. In addition to this evidence of individual learning, juveniles that observed tool use from human foster parents had an increased tool handling time and were influenced in their choice of object, demonstrating the influence of social learning on development. Kenward et al (2006) also found that subjects performed tool-oriented behaviour from 1 week after “branching” (defined as when subjects began to wander outside the nest; occurring at 25-26 days old), which increased in frequency each subsequent week. Across the developmental studies, tool use was most commonly measured as tool-oriented behaviour, via object manipulations and object choice (Fig. 2).

**Fig. 2.**
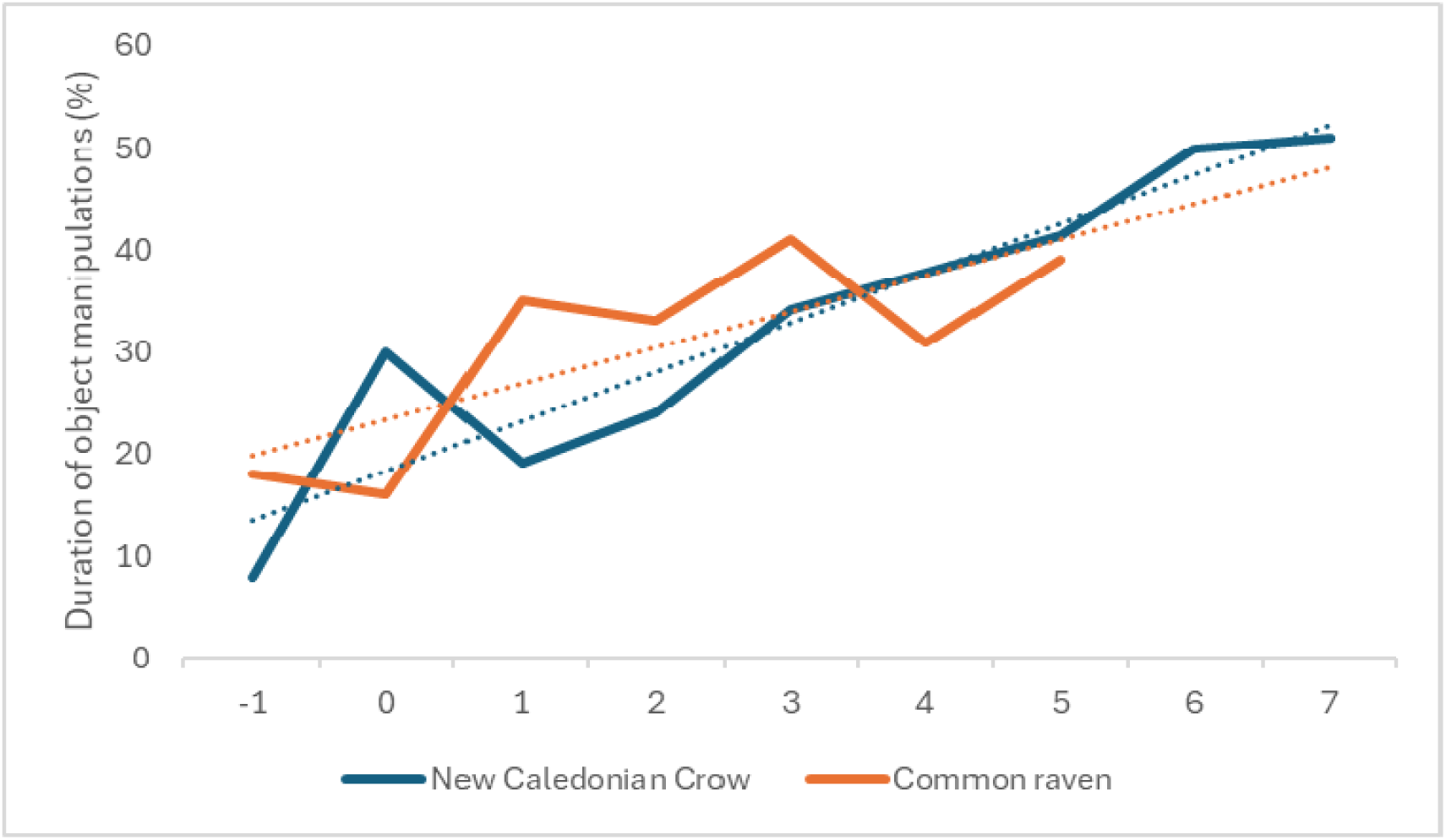
Duration of tool-oriented behaviours from observations of New Caledonian crows (Kenward et al, 2011; Kenward et al, 2006) and common ravens (Kenward et al, 2011). The x-axis represents fledging age of the birds (with -1 being 1 week pre-fledging). The New Caledonian crow percentages were averaged from the results of all tool-oriented behaviours (with the exception of precursor actions and functional insertions) in both Kenward et al (2011) and Kenward et al (2006), while the raven percentages were taken directly from the results of all tool-oriented behaviours (with the exception of precursor actions and functional insertions) in Kenward et al (2011). Precursor actions and functional insertions were not included due to a difference in types of measurement between papers, with Kenward et al, 2006 using percentage of time observed, while Kenward et al, 2011 used actions per minute

As indicated in Fig. 2, ravens and New Caledonian crows showed similar levels of object manipulations, which included evidence of simple action patterns that could be a precursor to tool use. Kenward et al (2011) suggested that the similarity of object manipulation duration was due to caching (ravens) and tool use (crows) requiring similar action patterns. However, tool use also requires object combinations, which New Caledonian crows performed significantly more of than ravens. Basic object combinations could be a precursor to tool use in corvids, and combinations such as probing and inserting can be considered direct tool-oriented behaviour, while caching is less likely to promote tool use (Auersperg et al, 2015). Auersperg et al (2015) found that ravens mainly performed caching as object combinations, while New Caledonian crows mainly performed inserting actions. Therefore, while caching may be an evolutionary precursor to tool use, tool-oriented behaviour may be a developmental precursor, hence why a habitual tool user (New Caledonian crow; Rutz & St Clair, 2012) may perform more of such object combinations than a non-habitual tool user (raven; Jacobs & Osvath, 2023).

#### Object manipulation and play

While object manipulations and combinations may be considered tool-oriented behaviour, when they are not overtly functional, they can also be a form of play (Auersperg et al, 2015). Animal play is typically defined as an apparent non-functional behaviour that is repeated and differs from other behaviours in terms of its context, its structure or occurs at an earlier age than when the behaviour becomes necessary. It is expressed when the animal is in a low stress or relaxed state (Burghardt, 2015). Play is often categorised into three forms: social play with conspecifics; locomotor-rotational play involving one’s own body, e.g. jumping; and physical play which includes object manipulation and combination (Osvath & Sima, 2014). Object play has been proposed to be a precursor to caching and tool use (O’Hara & Auesperg, 2017). Across juvenile to adult life stages, Auersperg et al (2015) found across three corvid species and three parrot species (and over life stages), object combinations during play correlated with problem-solving and physical cognitive abilities. In juvenile ravens, duration and frequencies of object manipulations were found to be consistent between age groups (3-6 months and 3-7 months respectively; Stöwe et al, 2006; Wenig et al, 2021). For reference, three months is past an early post-fledging age when ravens will already be regularly manipulating objects (see Figs. 2 and 3).

**Fig. 3.**
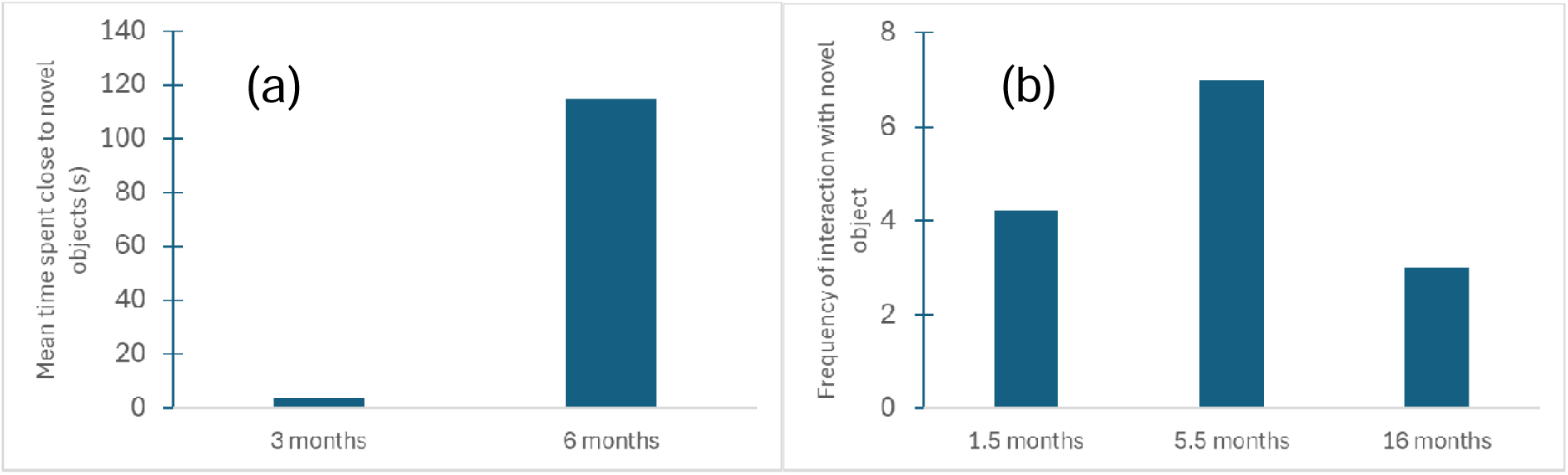
Interactions with novel objects across development in common ravens. Figure represents: (a) The median [respective] values of each object interacted with by 3-month-old and 6-month-old ravens were averaged (Stöwe et al, 2006) - referred to as the mean of time spent in close proximity (<1m) to the novel object. (b) The median [respective] values of frequency of interaction with all novel objects (not novel persons) from Miller et al (2015), with ages of the subjects averaged for each developmental stage (i.e. fledglings - 1-2 month = 1.5 months; juveniles - 3-8 months = 5.5 months; sub-adults - 14-18 months = 16 months)

Fig. 3 uses the species example of the common raven and presents object manipulation data from Stöwe et al (2006) - 3(a) and Miller et al (2015) - 3(b). These papers used different populations of captive ravens, but both found an increase in “novel object approach” – measured as time spent near novel objects in Stöwe et al (2006) and frequency of interactions in Miller et al (2015) - between birds aged 1.5-3 months old (fledging) to 5.5-6 months old (juvenile). These ages reflect when exploration may be important for ravens (O’Hara et al, 2017), therefore novel objects may be more willingly approached and interacted with (Miller et al, 2015). In Figure 3(b), once subjects reached 16 months (subadult), object manipulations significantly decreased. Adult ravens typically exhibit high levels of neophobia (Miller et al, 2022).

These trends in Fig. 3 - along with findings in Auersperg et al (2015) - indicate that object manipulation in ravens increases post-fledging, as exploratory behaviour appears particularly important for juveniles when interacting with new environments or stimuli (O’Hara et al, 2017) and decreases into sub-adulthood (Miller et al, 2015). Miller et al (2015) found the same object manipulation trend (frequency increases from fledging to juvenile; frequency decreases from juvenile to subadult) in carrion crows. We found that object manipulation was the main method of testing for evidence of developing physical cognition in young corvids, whether through tool-oriented behaviour, play-like object combinations or novel object manipulations.

#### Gaze following

Social cognition was more difficult to analyse comparatively than core or physical cognition, as outcome measures in this domain significantly varied across the literature. However, gaze following, one fundamental component of social cognition, was present in our coded output (Zeiträg & Osvath, 2022). Gaze following involves the ability to track the gaze direction of other individuals, such as humans, potential predators or conspecifics (Zeiträg & Osvath, 2022). Gaze following provides individuals with information about others’ attention, which can be useful for locating potential targets in the environment, like foraging opportunities or predators to be aware of (Zeiträg & Osvath, 2023). Developmentally, it offers a useful comparative framework, as it emerges early across many taxa and shows age-related changes in performance (Zeiträg et al, 2022). Despite methodological variation, developmental studies converge on the finding that gaze following emerges before maturity and reliably improves with age. A subject’s visual co-orientation with a conspecific or human demonstrator’s gaze is a primary paradigm. Within this paradigm, Zeiträg et al (2022) distinguishes between gaze following into the distance (GFD) and geometric gaze following (GGF), in which the subject must infer the target of gaze behind a visual barrier. GGF is considered cognitively more demanding, as it requires inference about another’s visual perspective. Ontogenetic studies support this distinction, with GFD emerging earlier across taxa and GGF developing later in the species that have been tested on both paradigms.

In corvids, common ravens provide a clear example of this pattern, as they were shown to follow a human experimenter’s gaze into distant space as fledglings, but not around barriers (GGF) until approximately six months of age (Bugnyar et al, 2004). Bugnyar et al (2004) and Schloegl et al (2008b), which found the same pattern in rooks, proposed that GFD and GGF differ in ecological relevance: scanning the sky may function as an anti-predator response that emerges early, whereas tracking gaze around barriers is more relevant to foraging and social competition and therefore develops later. Zeiträg et al (2022) also noted that the emergence of GGF coincides with the onset of hiding behind barriers during caching, suggesting a developmental milestone in understanding visual perspectives (Bugnyar et al, 2007).

Zeiträg & Osvath (2023) found that for young ravens, following the gaze of both human and conspecific demonstrators improved with age. Subjects were able to track gaze direction from behind a visual barrier 4 months after fledging, which corresponds with the age that ravens often become independent of their parents (Schloegl et al, 2007). Zeiträg and Osvath’s (2023) findings were that young ravens showed gaze following behaviour (“checking back”) from 30 days old and significantly more often with conspecific, than human, demonstrators. The tracking of gaze direction can aid in threat avoidance and aid in acquisition of other cognitive traits, such as theory of mind and social learning in human children (Zeiträg et al, 2022). Further research on corvid development of gaze following, particularly with conspecific demonstrators (Zeiträg & Osvath, 2023), can help us to better understand at which developmental stage different corvid species are likely to develop social traits in general.

### Objective 3: How does cognitive development vary across corvids and other taxa?

#### Comparison within the corvid family

16 corvid species were represented in the coded review output (S2 Table). 87% of species tested were from the *Corvinae* corvid subfamily, including common ravens and New Caledonian crows (6 papers included species within *Cyanocoracinae* or *Perisoreinae* subfamilies, including Florida scrub-jays and azure-winged magpies). Species tested varied in socio-ecological variables, including sociality, territoriality, caching and tool-use/ manufacture (Emery & Clayton, 2004), though few studies specifically tested the effects of socio-ecology on cognitive development. We outline some examples that include socio-ecological influences below.

#### Neophobia

Neophobia is influenced by age, though the developmental patterns vary by species. In Alalā (Hawaiian crow)*, Corvus hawaiiensis,* neophobia declines with age as juveniles are more neophobic than adults (Greggor et al, 2011). In other corvid species, like common ravens and carrion crows, it increases from juvenile to sub-adult life stages (Miller et al, 2015). In (primarily) adult corvids, neophobia is driven by differences in socio-ecology, as urban habitat use, adult sociality, maximum flock size and caching influenced object neophobia (Miller et al, 2022). More broadly, across 136 bird species from 25 taxonomic orders, bird species with specialist diets were more neophobic than those with generalist diets, and migratory species were more neophobic than nonmigratory species (ManyBirds Project et al, 2025)

#### Comparison with primates and other taxa

##### Why compare taxa?

It has been argued that primates and corvids have convergently evolved similar cognitive tools commonly associated with intelligence - such as causal reasoning, flexibility, imagination, and prospection - driven by comparable social and ecological pressures (Emery & Clayton, 2004). In addition to these similarities in adult cognitive performance, corvids and non-human primates also share aspects of their life histories that may be particularly relevant for cognitive development, including extended parental care, prolonged juvenile dependence and a comparatively long period before sexual maturity and dispersal. In primates, this extended juvenile period is thought to be critical for social learning and cultural transmission of valuable information like foraging strategies, tool use skills, and social norms (Whiten & van de Waal, 2018). Shared life-history traits and cognitive abilities motivate our comparisons of development across primates, corvids and parrots.

#### Similarities and differences in performance

##### Object permanence

Piagetian object permanence tasks have been applied developmentally across a wide range of taxa without excessive modification (see Objective 2 for examples). While there is variation in the maximum stage that different taxa reach, evidence suggests that object permanence advances universally over the course of development, with juveniles first succeeding on tasks involving partial occlusion before acquiring the ability to retrieve fully hidden objects and, later, objects undergoing visible and invisible displacement (Gómez, 2005). Across species, this advancement through stages appears sequential rather than abrupt, with individual variation in timing and completion of standardised tasks such as Uzgiris and Hunt’s Scale 1. For example, it would be unlikely for an individual to succeed at Stage 6 tasks before Stage 5 tasks, though within a stage it is not uncommon for individuals to succeed at tasks out of order.

In human children, object permanence develops early in life, with success on search-based tasks for fully occluded objects typically emerging around 8–12 months, although looking-time paradigms suggest sensitivity to object continuity at much younger ages (Baillargeon et al, 1985, 1987). In non-human primates, performance on comparable search and displacement tasks also improves across early development, with juveniles generally showing earlier success on visible displacement tasks than on tasks involving invisible displacement. While all great ape species tested reach Stage 6 (Barth & Call, 2006; Call, 2001; de Blois et al, 1998), some monkey species do not reliably master invisible displacements (Gómez, 2005). Common across all primates tested, however, is the tendency to make transitional A-not-B errors between Stages 5 and 6.

This developmental profile contrasts with those of domestic dogs and cats, for example, which develop object permanence relatively quickly, do not show A-not-B errors, but only reliably reach Stage 5 (Dumas & Doré, 1989; Gagnon & Doré, 1994; Gómez, 2005).

Developmental studies in corvids and parrots reveal a broadly similar progression to that observed in primates, including evidence of A-not-B errors, suggesting convergent developmental trajectories despite major differences in morphology and life history. Although humans likely progress more slowly than great apes, which progress more slowly than monkeys, direct comparisons of developmental speed are complicated by substantial differences in life history timing and length of juvenile periods.

#### Tool use

Tool use provides a complementary perspective on cognitive development across species because the acquisition of tool skills is mediated by social and ecological factors in addition to the progression of maturing cognitive and physical development. In addition to specific motor skills, tool users may show related understanding object concepts like causality, and executive function components such as planning and inhibitory control (Meulman et al, 2013). Spontaneous instances of tool use have been documented across a wide range of taxa, however we focus only on habitual tool users - species (or populations) where tool use is part of that population or group’s regular behavioural repertoire. Habitual tool use has been reported in multiple primate species (including chimpanzees, capuchins and macaques), several bird taxa and a small number of other mammals (reviewed in Meulman et al, 2013). While various corvid species have been able to manufacture and use tools as adults in laboratory settings (Bird & Emery, 2009a; see Giri & Garcia Pelegrin, 2025 for an overview), there are only two confirmed habitual tool users in the corvid family: the New Caledonian crow and the Hawaiian crow – the latter being functionally extinct in the wild (Rutz et al, 2016). In populations with habitual tool use, individuals often start exploring and manipulating materials as juveniles, and these early tool-related behaviours are characterised by systematic errors including incomplete action sequences, inappropriate materials, incorrect ordering of actions or application of otherwise correct actions toward the wrong substrates (Meulman et al, 2013).

Tool use development contrasts with other cognitive skills reviewed in this section (object permanence and gaze following), which are mastered early on in development, often around infancy. Across most habitual tool users, tool use is mastered relatively late in the juvenile period, depending on the complexity of the tool use ability (Meulman et al, 2013). For example, chimpanzees in the Goualougo Triangle appear to have a culture that involves the manufacturing of multiple types of tools to accomplish different aspects of the termite foraging process. Musgrave et al (2020) found that chimpanzees did not successfully complete the most complex tool task until an average age of over 11 years old - around the age that the species reaches sexual maturity. While simpler tool tasks can be accomplished at younger ages, this is still in contrast to the other cognitive skills reviewed in this paper, some of which can be achieved in infancy. Similar prolonged developmental trajectories have been reported in other habitual tool users, including but not limited to capuchin monkeys (*Cebus* sp.) (Falótico et al, 2024), long-tailed macaques (*Macaca fascicularis*) (Tan, 2017) and New Caledonian crows (Gray et al, 2010), with adult-like efficiency emerging only after months or years of refinement.

When controlling for differences in life history, it is not uncommon for more challenging tool tasks to be mastered close to the time of independence or sexual maturity across species. While gaze following and object permanence seem to mature internally, tool use, which is often both cognitively and physically demanding, is often thought to require more exposure (social learning) and repeated practice (Meulman et al, 2013). Research indicates that delayed acquisition cannot be fully explained by physical maturation - young individuals often possess sufficient strength or motor control to perform the relevant actions yet continue to show inefficient or inflexible performance (Meulman et al, 2013).

Laboratory studies of New Caledonian crows have shown that juveniles develop tool use similar to their wild counterparts even without a demonstrator, suggesting that genetic inheritance plays a significant role in these capabilities (Kenward et al, 2005). Studies on captive chimpanzees that show naïve individuals performing tool use behaviours seen in wild populations have raised questions of whether social learning is the primary means through which habitual tool users gain their skills (Bandini & Tennie, 2017). Kenward et al (2010) compared the ontogeny of caching in ravens to the ontogeny of tool use in New Caledonian crows, offering the hypothesis that the specialised tool behaviour in crows has evolutionary origins in caching (Fig. 3). Both New Caledonian crows and common ravens manipulated objects with similar frequencies throughout development (supported by Fig. 3), however, combinatorial manipulations, defined as placing an object in contact with another object or substrate, continued to increase in NC crows, but decreased in ravens 6 weeks after fledging. These findings support the proposal that there are phenotypic bias/predispositions in species that are habitual tool users, even in a laboratory setting without conspecific demonstrators (Kenward et al, 2010). Longitudinal research on taxa that are flexible tool users in the lab, but not in wild populations, could help further assess the extent to which habitual tool users are predisposed to tool use, the role of social learning in propagating tool use techniques, and also help elucidate the general developmental trajectory of problem-solving abilities.

#### Gaze following

Zeiträg et al (2022) review demonstrates that gaze following into the distance (GFD) is widespread across the animal kingdom, with evidence in fish and reptiles in addition to birds and mammals. On the other hand, geometric gaze following (GGF) seems to be an ancestral trait in the avian class, with evidence even in palaeognaths, but it has not been reported in fish or reptiles. Other key paradigms to test gaze following tend to fall under the object choice category, in which subjects must use gaze, posture, or gesture cues from an experimenter or conspecific to choose between objects. However, these have been less widely applied in developmental studies in corvids, but adult jackdaws (von Bayern & Emery, 2009) and Clark’s nutcrackers (Tornick et al, 2011) have performed successfully in this paradigm. As with every cognitive ability, different paradigms may be better suited to different species, so failure at one specific task does not necessarily indicate, in this case, poor social cognition or understanding of visual perspectives.

Dogs, for example, are able to follow cooperative human social cues without training (Hare et al, 2002), while many nonhuman primates are not (Zeiträg et al, 2022). This does not necessarily reflect poorer social cognition in primates, which often live in highly complex social environments. Rather, it likely reflects the competitive nature of primate foraging socioecology, and the generations of selective breeding for cooperation in domesticated animals (Hare & Tomasello, 2005). An additional example is that chimpanzees famously performed better in competitive versions of object choice and discrimination paradigms than traditional cooperative ones (Hare & Tomasello, 2004). In the wild, chimpanzees tend not to share monopolisable food resources, thus performing better at finding hidden food using an experimenter’s cues when the experimenter acted as a competitor as opposed to a cooperator (Hare & Tomasello, 2004).

In addition to socioecological factors, performance on gaze-following tasks is also shaped by morphology and experimental context. Species differ in the visibility of eye cues, reliance on head or body orientation, and attentional strategies, and across most taxa, subjects are generally more responsive to conspecifics than to human demonstrators (non-human primates, Kano & Call, 2014; ravens, Schaffer et al, 2020; ungulates, Zeitrag & Osvath, 2023). As with object permanence, these factors complicate the validity of direct comparisons of the same paradigms across taxa. In primates, including humans, individuals progress from responding to gross body orientation and gestures to later attending to more subtle eye movements (Gómez, 2005). In many species lacking a visible sclera, attention is more readily inferred from head or body orientation. In birds, gaze direction can be especially difficult to interpret due to lateral eye placement and the use of both monocular and binocular vision (Martin, 2007).

**Fig. 4** presents the percentage of subjects that performed gaze following at different early ages across species: common raven (Shloegl et al, 2007), rhesus macaque, *Macaca mulatta* (Tomasello et al, 2001) and wolf, *Canis lupus* (Range & Virányi, 2011). The measure reflects whether subjects look in a certain direction after seeing the experimenter look in the same direction, with both test and control trials. In control trials, experimenters would look either at or near the subject (Tomasello et al, 2001) and measure whether subjects looked in the same direction as in experimental trials. Fig. 4 suggests, from a descriptive perspective, that ravens and macaques may have perform gaze following at similar developmental ages. After this point, the macaques appear to show a consistent increase in this ability (of these three species) as they develop. The wolves seem to have some similar abilities at the same developmental stages as the macaques, however, their data did not follow a clear trend. The wolves in Range & Virányi (2011) had significant operant conditioning training during the developmental period, which may facilitate experimental participation. This could, however, have led them to develop an association of looking in the other direction when receiving a reward, even in control trials. After two failed control trials, the subjects may then have readjusted, explaining why most wolf subjects gaze followed at the first and last trial. All three species had a large number of individuals gaze following at their corresponding latest tested developmental stage.

**Fig. 4.**
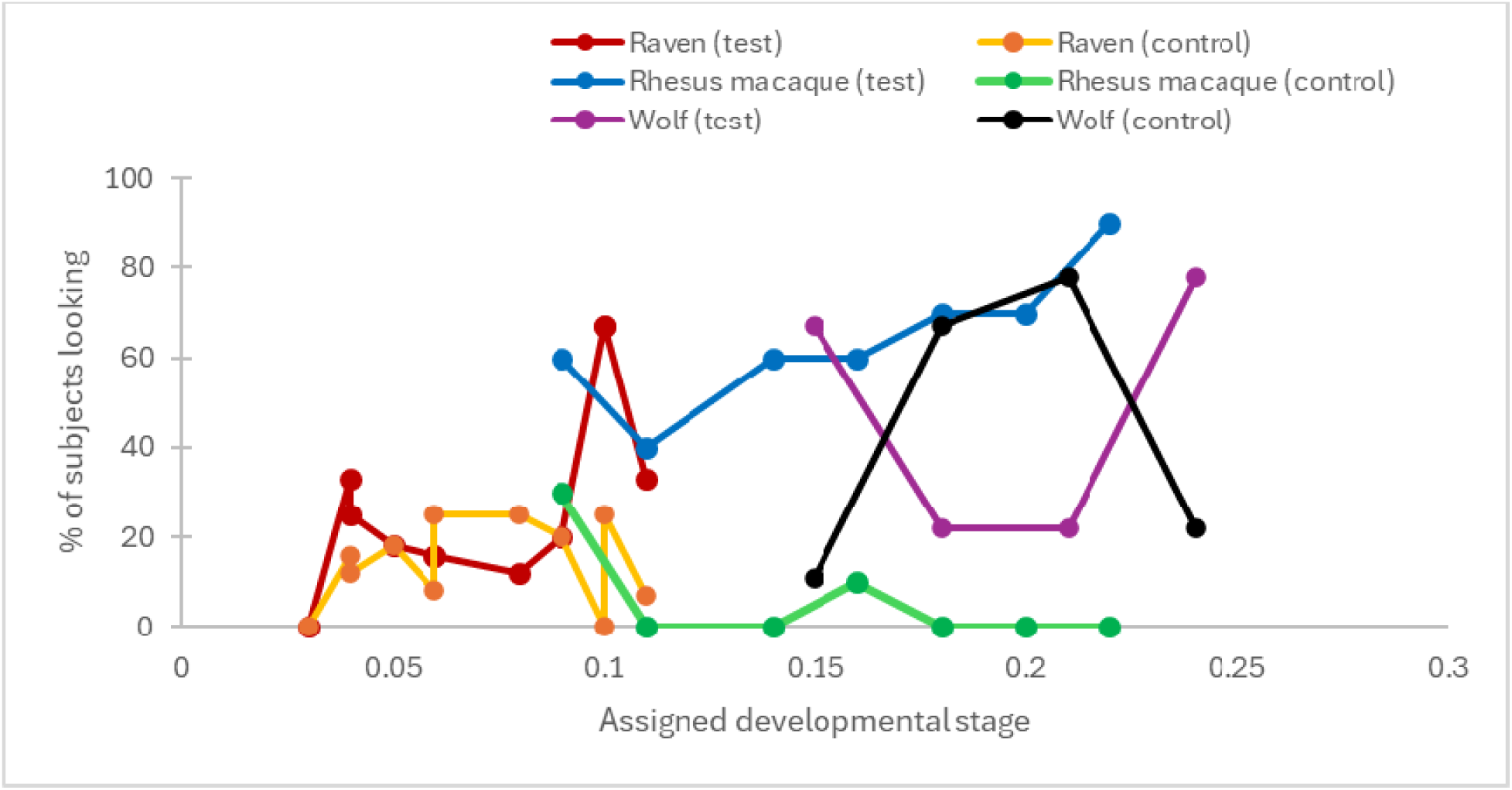
Comparison of the gaze following performances of three different species (common raven, rhesus macaque and wolf). The percentage of subjects that looked in the correct direction in test trials is reported, as well as the percentage that looked in the same direction in control trials, at each developmental age. The assigned developmental age is the age tested divided by the age of sexual maturity of the subjects (due to differences in lifespan and adulthood between species).

Listed sexual maturity for each species was: 3 years for ravens (Schloegl et al, 2007), 4 years for rhesus macaques (Tomasello et al, 2001) and 22 months for wolves - no age of adulthood mentioned in Range & Virányi (2011), instead this suggested age is taken from Mech et al (2016) and Müller et al (2025)

#### Embodied cognition

Another interesting aspect to consider in interspecies/ taxa comparisons is embodied cognition - the notion that the brain and body cannot be viewed as separate (Barton & Barrett, 2025; Wright & Clayton, 2025). Theories of embodied cognition argue that cognition is not a neutral process across species and is instead fundamentally intertwined with an organism’s body plan, perception and sensory environment (Foglia & Wilson, 2013). Traditional paradigms in comparative psychology have been influenced, and likely limited, by the use of human cognition as the standard baseline for comparison (Van Woerkum & Barrett, 2024). Tool use, for example, is highly dependent on an animal’s body plan, with primates’ possession of opposable thumbs being an obvious case. Birds, by contrast, rely primarily on the beak, head, and feet, with substantial variation across taxa. Parrots possess notably dexterous feet that can be brought to the beak during object manipulation, whereas many corvids exhibit morphological specializations related to food transport and caching, such as sublingual throat pouches. New Caledonian crows, the most prolific tool users within the corvid family, also show unique beak adaptations thought to support tool manufacture and control (Matsui et al, 2016). These anatomical differences suggest that similar cognitive capacities may be expressed through different motor solutions, and that differences in task performance may sometimes reflect physical constraints rather than cognitive ones.

Noting these differences in embodiment is particularly relevant given that many widely applied paradigms are adaptations of human experiments or theories originally developed for humans, such as Piagetian tasks. Even within paradigms initially designed for humans, researchers must therefore take motor development and physical differences into account when designing tasks. For example, researchers in the 1960s began exploring methods to measure object concept that require less motor coordination, finding evidence against the initial assertion put forth by Piaget that human infants attain Stage 4 object permanence at around nine months (Bower, 1967). When using violation-of-expectation paradigms that measure looking time rather than physically demanding object search tasks, for example, evidence for Stage 4 object permanence has been found in infants under five months of age (Baillargeon et al, 1985, 1987). Even body posture has been shown to impact infants’ success on the A-not-B task, with goal-directed reaching suggested to reflect an interaction between cognitive and physical processes rather than a failure of representation (Smith et al, 1999). If infants had formed a stable mental representation of the missing object, differences in posture would not be expected to affect task performance (Barton & Barrett, 2025).

All together, these examples serve as reminders that the outcomes of developmental studies may not directly reflect the timing of cognitive competence, highlighting the importance of considering embodied cognition when designing developmentally appropriate comparative tasks; the potential of magic-inspired experimental paradigms in comparative and developmental research is discussed further in the Future Directions section.

### Objective 4. How robust is the existing developmental evidence base?

#### Current issues within existing literature

Limitations within the existing literature include: relatively small sample sizes (min = 4; max = 224; median = 12; mean = 23.7 subjects per study), 74% tested in captive settings (12 field sites; 25 lab sites; 6 mixed lab-zoo or lab-field sites; 4 zoo sites), species biases and lack of age definition clarity (49% of papers coded with “partial” or “poor” age definition clarity, potentially impacting future replications). For wild compared to captive origin of subjects, 29% (16) of species tested had captive-bred subjects and 51% (28) had wild-bred subjects (20% were mixed or unclear). The field is currently lacking data on wild juveniles and on corvid species outside of those most commonly tested - of 139 corvid species currently recognised, only 16 species were represented in the reviewed literature, with 45% of identified articles on common raven (S2 Table). With regard to study design, 85% included repeated measures and 30% were longitudinal (14 papers longitudinal, 40 repeated measures, 8 cross-sectional), with some overlap in design across papers.

#### Current knowledge gaps

The major knowledge gaps appear to relate to cognitive domains/ abilities and related behaviours like executive function, causal reasoning, theory of mind, self-control, metacognition, communication, cooperation, mental time travel, spatial learning and social learning. There are established paradigms that have been adapted from the human/ child development literature, which could be implemented to assess corvid development. For example, the ability to delay gratification, i.e. to obtain a more valuable outcome in the future, through tolerating a delay or investing a greater effort in the present (Beran et al, 2016), which is a measure of self-control, has been demonstrated in (primarily adult) Eurasian jays, New Caledonian crows, common ravens, carrion crows, California scrub-jays, blue jays and pinyon jays (Clayton et al, 2005; Dufour et al, 2012; Hillemann et al, 2014; Miller et al, 2019; Stephens & Anderson, 2001; Stephens & Dunlop, 2009). Using the “rotating-tray” paradigm, Eurasian jays were found to flexibly switch their choices for a delayed, preferred reward over an immediate, less-preferred reward in the presence of competitors, while New Caledonian crows selected the delayed, preferred reward irrespective of social presence (Miller et al, 2023). However, we do not yet know how this ability develops or changes across ontogeny and there is scope to further assess the flexibility of self-control across contexts.

#### Limitations within our review

Limitations within our review include: potential of missed key literature, despite the systematic review approach - we selected a restricted time period for focus (20 years) with the expectation that it would be possible to code papers from this period as comparably as possible for all coded variables. Our key word searches also only applied to titles and abstracts, so literature with juvenile subjects could have been missed if age classes were only mentioned within the article, but not the title or abstract. We found there was some overlap of “cognition” with “behaviour”, which was difficult to tease apart using search term keywords, therefore, we conducted two searches (with and without “cognition” terms included), then combined findings. We then included all articles for coding that were related to corvids and development/ ontogeny, whether they specified the focus as “cognition” or more so on “behaviour” (for example, anti-predator behaviour; social foraging behaviour).

## 4. Future Directions

Key areas for future research include addressing the knowledge gaps outlined above and answering broader questions like how early cognitive abilities relate to later behavioural or fitness-related outcomes in corvids and other species, such as survival, dispersal, reproductive success and social integration. More standardised comparative approaches would allow for teasing apart internal and external influences on development. For example, in Miller et al (2015; 2016; 2025), we tested object manipulation, exploration and caching behaviour across development, from fledging to sub-adult stages, and varying social contexts in identically reared, housed and tested common ravens and carrion crows. Although these two species are closely related, we identified both converging and diverging patterns of behaviour over development, likely relating to broader socio-ecological species differences, such as habitat use and caching proficiency.

### Repeatability

Repeatability estimates may vary depending on a range of factors, including taxa, sex, age, testing setting (captive vs wild), number of measures and the interval between measures, for example, being higher when observation intervals were short (Bell et al, 2009). Only 13% (6) papers, across 5 species, in our sample specifically tested for individual repeatability. It would be beneficial to understand how stable or repeatable individual differences in cognition are from fledging/juvenile to adulthood life stages, including potential dependence on physical and social development and environments (Damini et al, 2025). Neophobia was found to be largely repeatable across (primarily adult) bird individuals and species, including corvids (ManyBirds Project et al, 2025; Miller et al, 2022). While in ravens and crows over ontogeny, individuals were highly inconsistent - or flexible - in exploration behaviour, though in the ravens only, conspecific presence promoted behavioural similarities (Miller et al, 2016). Similarly, in four other (primarily adult) corvid species, individual exploration responses were not repeatable (Vernouillet & Kelly, 2020). We recognise that assessing repeatability over development requires longitudinal testing, which brings logistical considerations, including funding pressures.

### Science of Magic

An exciting area for future research is the integration of magic-inspired violation of expectancy (VoE) paradigms, as controlled probes into cognitive understanding and development. In human infants, classic VOE paradigms demonstrate that seemingly “impossible” physical outcomes reliably increase looking time relative to matched “possible” outcomes, probing object permanence, causal reasoning and other cognitive abilities (Baillargeon et al, 1985; Garcia-Pelegrin et al, 2024). VoE looking-time paradigm was used to test the ontogeny of support relations in Eurasian jays, controlling for experience of different support types, finding that by 9 months old, jays looked more at a tool moving an item that was not in contact than one that was in contact (i.e. impossible outcome; Davidson et al, 2017). Garcia-Pelegrin et al (2023) found that manual action expectations are constrained by biomechanical ability in three non-human primate species, illustrating how sleight of hand magic effects can be used to comparatively test predictive cognition. Furthermore, Eurasian jays were found to respond in a similar vein to humans in that they were fooled by fast sleight-of-hand effects that rely on rapid object transfer, but contrasted in that they were not misled by the “French drop” illusion, which depends on preconceived expectations about hand biomechanics that even birds with extensive experience being fed by human hands do not have (Garcia-Pelegrin et al, 2021).

These findings indicate that perceptual biases revealed by magic illusions are not unique to humans, indicating shared cognitive constraints across diverse species, including monkeys and corvids.

Smith et al (2025) found that kea, *Nestor notabilis*, though not cockatoos, *Cacatua goffiniana*, were susceptible to a “bait and switch” magic trick. Using infrared thermography, the study found no evidence for the effect of violations of expectation on subject periorbital temperature - a measurement of physiological response (Smith et al, 2025). The next steps may include exploring magic-inspired paradigms from a developmental perspective in corvids and other species, concurrently assessing relevant physiological, anatomical and behavioural measures, such as pupil dilation, emotional responses, facial expression and neurological changes (Smith et al, 2025), as well as measures of surprise.

### Theory of Mind

Additional research on the ontogeny of caching would likely be an ecologically sound avenue for investigating the development of theory of mind in corvids. As mentioned earlier, although adult corvids show sophisticated cache protection strategies that are often interpreted as evidence for theory of mind, comparatively little is known about how these abilities emerge developmentally. In adult corvids, experience projection has been shown to influence cache protection behaviour. In an experiment with adult Florida scrub-jays, individuals with prior experience pilfering from others’ caches were significantly more likely to re-cache food after being observed caching by conspecifics, suggesting that cache protection strategies may depend on relating one’s own past experience as a pilferer to the risk of future theft from another individual (Emery & Clayton, 2001).

In the existing research on caching ontogeny though, it remains difficult to determine whether early instances of cache protection reflect experience-based learning (e.g., greater cache recovery success when caching alone than when observed) or a more explicit understanding of others as individuals with different perspectives and access to information. This distinction reflects broader challenges in the field of comparative cognition, where similar behavioural outcomes may arise from different underlying mechanisms. More research using developmental approaches could help address this ambiguity by tracking when cache protection behaviours and deceptive tactics first appear, how they change with age and experience, and whether this timing aligns with improvements in related skills such as gaze following or inhibitory control. In parallel, developmental studies of gaze following, particularly geometric gaze following, may provide additional insight into the emergence of visual perspective taking, as this ability is thought to require inference about what another individual can see (Zeiträg et al, 2022).

Magic-based paradigms may also provide a useful framework for Theory of Mind research, as caching corvids take advantage of others’ attentional blind spots and employ deceptive tactics – similar to what magicians do (Garcia-Pelegrin et al, 2024). Examining the development of sensitivity to sleight of hand techniques, and the timing of deceptive tactics emerging in corvids may also help distinguish when birds begin taking the perspectives of others into consideration. Overall, longitudinal studies that combine caching paradigms with controlled experiments of visual perspective taking may be especially valuable for clarifying how and when social cognition develops in species with sophisticated caching behaviours.

## 5. Conclusion

Overall, this review synthesises the breadth of research on cognitive development in corvids from the last two decades. Across the most commonly researched domains, we found that cognitive development is generally characterised by gradual improvements influenced by both internal maturation and/or experience. Foundational abilities such as object permanence and gaze following appear relatively early in development, while behaviours like tool use and socially informed caching techniques emerge later on, indicating the possible influence of extended practice or learning. While considering variation in timing and expression, cross-species comparisons also suggest broadly similar developmental patterns across taxa, though also highlight the need to consider differences in life history, morphology and socioecological factors. Nonetheless, the existing developmental evidence base in corvids remains limited by small sample sizes, species biases, inconsistent age definitions and a strong reliance on captive populations, with juvenile wild corvids and most corvid species largely unrepresented in the reviewed literature. Major knowledge gaps remain for topics such as executive function, causal reasoning, self-control, theory of mind and social learning, particularly from an ontogenetic perspective. In order to address existing knowledge gaps in these domains, future research should embrace longitudinal approaches and creative experimental designs which take ecology and embodied cognition into consideration.

## Acknowledgements

Thank you to Dr James Davies, University of Bristol, for related discussions and his role in co-writing the awarded Leverhulme Trust Research Grant, which funded RM’s position (Oct 2025-Feb 2026).

## Author Contributions

RM conceived the review focus, research objectives and designed the full outline. RM and EC designed the search terms and conducted the systematic review searches, with EC filtering, combining and cross-checking the outputs. RM, EC and AT coded the articles. RM, EC and AT wrote the first full draft of the manuscript, and NSC provided editing and feedback on the drafts. RM and NSC provided supervision to EC and AT. NSC provided overall direction and funding, which funded RM’s position.

## Data Availability

The data (including full coding output) is available via Figshare: 10.6084/m9.figshare.31410981 (private link: https://figshare.com/s/b88ae401b55b99b35d57).

## Supplementary Information Legends

**Text S1:** Coding definitions for systematic review

**Table S1:** Overview of cognitive domains tested in relation to development/ ontogeny within corvids.

**Table S2:** Overview of abilities tested in relation to development/ ontogeny within corvids.

**Table S3:** Papers from coded review output with corresponding tested species, specific cognitive abilities tested, developmental trajectories, outcome measures and environmental effects.

**Table S4:** Uzgiris & Hunt Scale 1 task / Object permanence development across corvids and parrots

**Table S5:** Taxonomic distribution of developmental studies on object permanence, gaze following and tool use

