## Supplementary Information for "Development of cognition in corvids"

**Table S1***. Overview of cognitive domains tested in relation to development/ ontogeny within corvids*. Core/ foundational domain = object permanence, caching, learning, motor self-regulation, perception. Physical cognition = casual reasoning, tool use/ manufacture. Social cognition = social learning, gaze following, social behaviour, communication. Physical/social cognition = object manipulation, exploration, neophobia and play

| **Corvid species** | **No. of papers** | **“Core”**  **cognition** | **“Social” cognition** | **“Physical” cognition** | **References** |
| --- | --- | --- | --- | --- | --- |
|  | **Total % and no. of species tested per domain** | | | |  |
|  | 16 | 69% (11 species) | 56% (9 species) | 31% (5 species) |  |
| American crow, *Corvus brachyhyrhynchos* | 1 |  | 1 |  | Mates et al, 2014 |
| Azure-winged magpie, *Cyanopica cyanus* | 2 | 1 | 1 |  | Wang et al, 2021; Wang et al, 2023 |
| Black-billed magpie, *Pica hudsonia* | 1 | 1 |  |  | Clary et al, 2014 |
| Californian scrub-jay, *Aphelocoma californica* | 1 | 1 |  |  | Salwiczek et al, 2009 |
| Clark’s nutcracker, *Nucifraga columbiana* | 1 | 1 |  |  | Clary et al, 2014 |
| Canada jay, *Perisoreus canadensis* | 1 |  | 1 |  | Freeman et al, 2021 |
| Eurasian jay, *Garrulus glandarius* | 2 | 1 |  | 1 | Davidson et al, 2017; Zucca et al, 2007 |
| Common raven, *Corvus corax* | 21 | 10 | 16 | 1 | Auersperg et al, 2015; Boucherie et al, 2020; Beck et al, 2020; Braun et al, 2012; Bugnyar et al, 2007; Damini et al, 2025; Gallego-Abenza et al, 2020; Gallego-Abenza et al, 2022; Gallup et al, 2022; Jacobs et al, 2019; Kabadayi et al, 2017; Kenward et al, 2011; Loretto et al, 2020; Miller et al, 2015, 2016, 2025; Scheid and Bugnyar, 2008; Schloegl et al, 2007; Stowe et al, 2006; Wenig et al, 2021; Zeiträg and Osvath, 2023 |
| Carrion crow, *Corvus corone* | 5 | 4 | 3 |  | Hoffman et al, 2011; Miller et al, 2015; 2016; 2025; Wascher et al, 2025 |
| Western Jackdaw,  *Coloeus monedula* | 6 | 3 | 4 |  | Auersperg et al, 2015; Schwab et al, 2008; Scheid et al, 2008; Ujfalussy et al, 2013; von Bayern et al, 2007; Zandberg et al, 2014 |
| New Caledonian crow, *Corvus* *moneduloides* | 8 | 1 | 3 | 6 | Auersperg et al, 2015; Holzhaider et al, 2010a; 2010b; 2011; Hunt et al, 2007; Kenward et al, 2006; 2005; 2011 |
| Florida scrub-jay, *Aphelocoma coerulescens* | 2 | 2 | 1 |  | Bebus et al, 2016; Fuirst et al, 2020 |
| Hawaiian crow/ ʻAlalā, *Corvus hawaiiensis* | 1 |  |  | 1 | Greggor et al, 2020 |
| Large-billed crow, *Corvus macrorhynchos* | 1 |  | 1 |  | Obozova et al, 2018 |
| Pinyon jay, *Gymnorhinus cyanocephalus* | 1 | 1 |  |  | Stafford et al, 2006 |
| Rook, *Corvus frugilegus* | 1 |  |  | 2 | Tebbich et al, 2007 |

**Table S2***. Overview of abilities tested in relation to development/ ontogeny within corvids*. * = social behaviour includes anti-predator behaviour, social structure, social foraging, yawning. + = learning includes spatial, associative and reversal learning

| **Corvid species** | **No. of papers** | **Object permanence** | **Caching** | **+ Learning** | **Motor self-regulation** | **Perception** | **Neophobia/ exploration** | **Social Learning** | **Object manipulation and play** | **Gaze Following** | *** Social Behaviour** | **Communication** | **Causal reasoning** | **Tool use/ manufacture** | **References** |
| --- | --- | --- | --- | --- | --- | --- | --- | --- | --- | --- | --- | --- | --- | --- | --- |
| American crow, *Corvus brachyhyrhynchos* | 1 |  |  |  |  |  |  |  |  |  |  | 1 |  |  | Mates et al, 2014 |
| Azure-winged magpie, *Cyanopica cyanus* | 2 | 1 |  |  |  |  |  |  |  |  |  | 1 |  |  | Wang et al, 2021; Wang et al, 2023 |
| Black-billed magpie, *Pica hudsonia* | 1 |  |  |  |  | 1 |  |  |  |  |  |  |  |  | Clary et al, 2014 |
| Californian scrub-jay, *Aphelocoma californica* | 1 | 1 | 1 |  |  |  |  |  |  |  |  |  |  |  | Salwicxek et al, 2009 |
| Clark’s nutcracker, *Nucifraga columbiana* | 1 |  |  |  |  | 1 |  |  |  |  |  |  |  |  | Clary et al, 2014 |
| Canada jay, *Perisoreus canadensis* | 1 |  |  |  |  |  |  |  |  |  | 1 |  |  |  | Freeman et al, 2021 |
| Eurasian jay, *Garrulus glandarius* | 2 | 1 |  |  |  |  |  |  |  |  |  |  | 1 |  | Davidson et al, 2017; Zucca et al, 2007 |
| Common raven, Corvus corax | 21 | 1 | 7 | 1 | 1 |  | 3 | 1 | 4 | 2 | 5 | 1 |  | 1 | Auersperg et al, 2015; Boucherie et al, 2020; Beck et al, 2020; Braun et al, 2012; Bugnyar et al, 2007; Damini et al, 2025; Gallego-Abenza et al, 2020; Gallup et al, 2022; Kabadayi et al, 2017; Miller et al, 2015, 2016, 2025; Schloegl et al, 2007; Stowe et al, 2006; Wenig et al, 2021; Zeiträg and Osvath, 2023; Jacobs et al, 2019; Kenward et al, 2011; Scheid and Bugnyar, 2008; Gallego-Abenza et al, 2022; Loretto et al, 2020 |
| Carrion crow, Corvus corone | 5 | 1 | 3 |  |  |  | 3 |  | 1 |  |  | 1 |  |  | Hoffman et al, 2011; Miller et al, 2015; 2016; 2025; Wascher et al, 2025 |
| Western Jackdaw, Coloeus  monedula | 6 | 1 | 1 | 1 |  |  |  | 1 | 2 |  | 1 | 1 |  |  | Auersperg et al, 2015; Schwab et al, 2008; Scheid et al, 2008; Ujfalussy et al, 2012; von Bayern et al, 2007; Zandberg et al, 2014 |
| New Caledonian crow, Corvus moneduloides | 8 |  |  | 1 |  |  |  | 1 | 1 |  | 1 |  |  | 6 | Auersperg et al, 2015; Holzhaider et al, 2010a; 2010b; 2011; Hunt et al, 2007; Kenward et al, 2006; 2005; 2011 |
| Florida scrub-jay, Aphelocoma coerulescens | 2 |  | 1 | 1 |  |  | 1 |  |  |  |  |  |  |  | Bebus et al, 2016; Fuirst et al, 2020 |
| Hawaiian crow/ ʻAlalā, Corvus hawaiiensis | 1 |  |  |  |  |  | 1 |  |  |  |  |  |  |  | Greggor et al, 2020 |
| Large-billed crow, Corvus macrorhynchos | 1 |  |  |  |  |  |  |  |  |  | 1 |  |  |  | Obozova et al, 2018 |
| Pinyon jay, Gymnorhinus cyanocephalus | 1 |  |  | 1 |  |  |  |  |  |  |  |  |  |  | Stafford et al, 2006 |
| Rook, Corvus frugilegus | 1 |  |  |  |  |  |  |  |  |  |  |  | 1 | 1 | Tebbich et al, 2007 |

**Table S3***. Papers from coded review output with corresponding tested species, specific cognitive abilities tested, developmental trajectories, outcome measures and environmental effects.* Core/ foundational domain = object permanence, caching, learning, motor self-regulation, perception. Physical cognition = casual reasoning, tool use/ manufacture. Social cognition = social learning, gaze following, social behaviour, communication. Physical and social cognition = object manipulation, exploration, neophobia and play. Social behaviour includes anti-predator behaviour, social structure, social foraging, yawning. Learning includes spatial, associative and reversal learning

| **Paper** | **Species** | **Cognitive domain** | **Specific ability tested** | **Life stages tested** | **Developmental trajectory** | **Outcome measures** |
| --- | --- | --- | --- | --- | --- | --- |
| Holzhaider et al, 2010a | New Caledonian crow | Physical | Tool use | Juvenile, adult | Improves with age | Object manipulation duration |
| Holzhaider et al, 2010b | New Caledonian crow | Physical | Tool use/ manufacture | Juvenile | Improves with age | Frequency of tool use events |
| Hunt et al, 2007 | New Caledonian crow | Physical | Tool use | Fledging, adult | Not analysed | Trials to criterion |
| Kenward et al, 2011 | New Caledonian crow, Common raven | Physical | Tool use | Fledging, juvenile | Improves with age | Object manipulation duration |
| Kenward et al, 2005 | New Caledonian crow | Physical | Tool use | Juvenile | Not analysed | Number of times tool used in attempt of food retrieval |
| Tebbich et al, 2007 | Rook | Physical | Tool use | Juvenile | Not analysed | Trials to criterion |
| Kenward et al, 2006 | New Caledonian crow | Physical | Tool use | Fledging, juvenile | Improves with age | Object manipulation duration, object choice |
| Von bayern et al, 2007 | Jackdaw | Social, physical | Food/object sharing | juvenile | Declines with age | Foraging behaviour |
| Miller et al, 2025 | Common raven,  Carrion / hooded crow | Social, physical, core | Exploration, caching | Fledging, juvenile, subadult | Improves with age | Frequency of object interaction |
| Miller et al, 2015 | Common raven,  Carrion / hooded crow | Social, physical, core | Exploration, caching | Fledging, juvenile, subadult | Declines with age | Frequency of object interaction |
| Miller et al, 2016 | Common raven,  Carrion / hooded crow | Social, physical, core | Exploration, caching | fledging, juvenile, subadult | No pattern | Frequency of object interaction, caching frequency |
| Stöwe et al, 2006 | Common raven | Social, physical | Exploration, object manipulation | Juvenile | Stable | Object manipulation duration, frequency of object interaction-s |
| Beck et al, 2020 | Common raven | Core | Caching | Juvenile | No pattern | Frequency of caching trips, cache probability, cache distance |
| Fuirst et al, 2020 | Florida scrub jay | Core | Caching | First year, second year, experience-d adult | Stable | Condition of cache sites, position of cache sites |
| Auersperg et al, 2015 | Common raven, jackdaw, New Caledonian crow (other species: red-shouldered macaw, black-headed caique, Goffin’s cockatoo) | Social, physical | Object manipulation, play | Juvenile, subadult-juvenile, adult | Not analysed | Frequency of object combinations |
| Wenig et al, 2021 | Common raven | Social, physical | Play | Juvenile | Stable | Object manipulation duration, frequency of object manipulat-ions |
| Damini et al, 2025 | Common raven | Social | Anti-predator responses | Mixed ages | improves with age | General behaviour |
| Schwab et al, 2008 | Jackdaw | Social | Social learning | Fledging, juvenile, subadult | Improves with age | Affiliation behaviour, object manipulation duration |
| Freeman et al, 2021 | Canada jay | Social | Social status | Nestling, juvenile | Stable | Physiology |
| Wang et al, 2023 | Azure-winged magpie | Social | Communication, vocal repertoire | Nestling, juvenile, adult | Stable | General behaviour |
| Gallup et al, 2022 | Raven | Social | Contagious yawning | Juvenile | No pattern | General behaviour |
| Braun et al, 2012 | Common raven | Social | Social grouping | Juvenile, subadult, adult | Not analysed | General behaviour |
| Obozova et al, 2018 | Large-billed crow | Social | Social interactions | Fledging then juvenile | U-shaped | General behaviour |
| Schloegl et al, 2007 | Common raven | Social | Gaze following | Nestling, fledging | Improves with age | Gaze following |
| Zeiträg and Osvath, 2023 | Common raven | Social | Gaze following | Fledging, juvenile | Improves with age | Gaze following |
| Holzhaider et al, 2011 | New Caledonian crow | Social | Social structure | Juvenile, adult | Not analysed | General behaviour inc. foraging behaviour |
| Wascher and Youngblood, 2025 | Carrion /hooded crow | Social | Communication | Juvenile, adult | Declines with age | Call sequences in response to general behaviour |
| Boucherie et al, 2020 | Common raven | Social | Social behaviour | Juvenile | Not analysed | General behaviour and frequency of object manipulations |
| Gallego‐Abenza et al, 2020 | Common raven | Social | Social foraging behaviour | Juvenile, subadult, adult | Improves with age | Foraging behaviour |
| Gallego-Abenza et al, 2022 | Common raven | Social | Social attention/recognition | Juvenile | Not analysed | General behaviour in response to calls |
| Loretto et al, 2020 | Common raven | Social | Social learning | Juvenile | Not analysed | Trials to criterion, type of contact with box |
| Mates et al, 2014 | American crow | Social | Communication | Subadult, adult | No pattern | Foraging behaviour in response to calls, general behaviour in response to calls |
| Zandberg et al, 2014 | Jackdaw | Social | Call recognition/discrimination | Nestling | Improves with age | General behaviour in response to calls |
| Bugnyar et al, 2007 | Common raven | Core | Caching, object permanence | Fledgling, juvenile | Improves with age | General behaviour |

**Table S4***. Uzgiris & Hunt Scale 1 task / Object permanence development across corvids and parrots.* *Age at which last task was completed (so in task 5-9, age at which 9 was completed). Azure-winged magpies excluded because even though subjects were reported to be juveniles, study did not include developmental timing

| **Species** | **Estimated fledging age**  **(days)** | **Stage 3**  **Task 3** | **Stage 4**  **Task 4** | **Stage 5**  **Tasks**  **5-9** | **Stage 6**  **Tasks 10-14** | **Stage 6**  **Task 15** | **“A-not-b” errors** | **Reference** |
| --- | --- | --- | --- | --- | --- | --- | --- | --- |
| California Scrub-jay | 19 | g2: 28  g1: 32 | 35 | Not tested | Not tested | Not tested | Not tested | Salwiczek et al, 2009 |
| Eurasian jay  (mean age; younger jays) | 20 | 31 | 37 | 41 | 47 | 47 | No errors | Zucca et al, 2007 |
| Eurasian magpie  (mean age) | 25 | 32 | 57 | 107 | 180 | Did not pass | No errors | Pollok et al, 2000 |
| Common raven  (median ages, estimate based on weeks postfledging, two groups at different facilities) | 40 | Uvm: 47  Klf: 47 | Uvm: 54  Klf: 54 | Uvm:  89  Klf: 89 | Uvm: 110  Klf: 110 | Uvm: did not pass  Klf: 110 | Errors present | Bugnyar et al, 2007 |
| Carrion crow  (median ages) | 32 | 37 | 42 | 67 | 71 | Did not pass | 5/5 subjects made errors | Hoffmann et al, 2012 |
| Jackdaw  (mean ages) | 26-31 | 36.6 | 41 | 61 | 74.3 | 80.7 | 2/8 subjects made errors | Ujfalussy et al, 2012 |
| Grey parrot  (n=1) | 70-84 | 63 days | 112 days | 140 days | 154 days | 147 days* | Errors present | Pepperberg et al, 1997 |
| Kakariki  (Parent-raised vs. hand-raised groups) | 35-49 | PR: 63 days  HR: 56 days | PR: 108 days  HR: 73 days | PR: 150 days  HR: 133 days | PR: 164 days  HR: 161 days | PR: 164 days  HR: 161 days | Errors present | Funk, 1996 |

**Table S5***. Taxonomic distribution of developmental studies on object permanence, gaze following and tool use in non-human animals.* For the cross-species comparison table, we did not conduct a systematic literature review. Instead, we used existing reviews of object permanence, gaze following and tool use to identify representative studies across taxa and then screened these sources for papers that focused on ontogenetic development or explicitly mentioned using multiple juvenile subjects (Gómez, 2005; Jaakkola, 2014; Meulman et al, 2013; Zeiträg et al, 2022). Citations provided for each species are thus illustrative rather than exhaustive – for example, although numerous studies examine chimpanzee tool-use ontogeny in different populations, a single representative example was included in this table.

| **Species tested *developmentally* or as juveniles** | **Object permanence** | **Gaze following** | **Tool use (habitual)** |
| --- | --- | --- | --- |
| Corvids | Eurasian magpie (Pollok et al, 2000),  Eurasian jay (Zucca et al, 2007).  Common raven (Bugnyar et al, 2007),  Carrion crow (Hoffmann et al, 2012),  Jackdaw (Ujfalussy et al, 2012),  California scrub-jay (Salwiczek et al, 2009),  Azure-winged magpie (Wang et al, 2021) | Common raven  Rook | New Caledonian crow (Kenward et al, 2006) |
| Parrots | Grey Parrot *Psittacus erithacus* (Pepperberg et al, 1997),  Kakariki *Cyanoramphus auriceps* (Funk, 1996) | [Grey parrot](https://link.springer.com/article/10.1007/s10071-008-0163-2) (Giret et al, 2008) |  |
| Other birds | [Domestic chicken](https://elifesciences.org/articles/67208) *Gallus gallus domesticus* (Szabó et al, 2022) | [Greylag goose](https://link.springer.com/article/10.1007/s10071-011-0381-x), *Anser anser* (Kehmeier et al, 2011),  [Bobwhite quail](https://pmc.ncbi.nlm.nih.gov/articles/PMC2764337/), *Colinus virginianus* (Jaime et al, 2009),  [Domestic chicken](https://www.sciencedirect.com/science/article/pii/S0166432806006231), *Gallus gallus domesticus* (Rosa Salva et al, 2007) | [Woodpecker finch](https://royalsocietypublishing.org/rspb/article-abstract/268/1482/2189/71004/Do-woodpecker-finches-acquire-tool-use-by-social?redirectedFrom=fulltext), *Camarhynchus pallidus* (Tebbich et al, 2001),  [Green-backed heron](https://onlinelibrary.wiley.com/doi/abs/10.1111/j.1474-919X.1986.tb02677.x) *Butorides sp*. (Higuchi, 2008) |
| Great Apes | Chimpanzee, *Pan troglodytes* (Wood et al, 1980),  [Orangutan](https://www.eva.mpg.de/documents/AmericanPsychologicalAss/Call_Object_JCompPsych_2002_1556106.pdf), *Pongo pygmaeus* (Call, 2001),  [Gorilla](https://www.taylorfrancis.com/chapters/edit/10.4324/9781315808055-5/early-sensorimotor-development-gorilla-giovanna-spinozzi-francesco-natale), *Gorilla gorilla* (Spinozzi & Natale, 2019) | [Chimpanzee](https://psycnet.apa.org/record/2001-00389-007), (Tomasello et al, 2001),  [Bonobo](https://research.ebsco.com/c/dvoqdt/search/details/5ceam423of), *Pan paniscus* (Bräuer et al, 2005),  [Gorilla](https://research.ebsco.com/c/dvoqdt/search/details/5ceam423of) (Bräuer et al, 2005),  [Orangutan](https://research.ebsco.com/c/dvoqdt/search/details/5ceam423of) (Bräuer et al, 2005) | [Chimpanzee](https://onlinelibrary.wiley.com/doi/abs/10.1002/ajpa.24125) (Musgrave et al, 2020), [Orangutan](https://www.cambridge.org/core/journals/evolutionary-human-sciences/article/ontogeny-%5B%E2%80%A6%5Dehaviour-in-wild-orangutans/81CE0FC52D716D58A20BC08AFC0D5752) (Schuppli et al, 2021) |
| Other primate | [Japanese Macaque](https://www.researchgate.net/publication/16013358_Cognitive_development_in_a_Japanese_macaque_Macaca_fuscata), *Macaca fuscata* (Antinucci et al, 1982),  [Rhesus macaque](https://www.researchgate.net/publication/232431138_Piagetian_object_permanence_in_the_infant_rhesus_monkey), *Macaca mulatta* (Wise et al, 1974)*,*  Crab-eating macaque *Macaca fascicularis* (Schino et al, 1990),  Tufted capuchin *Cebus apella* (Schino et al, 1990) | [Southern pig-tailed macaque,](https://pubmed.ncbi.nlm.nih.gov/11095722/) *Macaca nemestrina* (Ferrari et al, 2000),  [Rhesus macaque,](https://psycnet.apa.org/record/2001-00389-007) *Macaca mulatta* (Tomasello et al, 2001),  [Barbary macaque,](https://onlinelibrary.wiley.com/doi/10.1111/j.1467-7687.2010.00956.x) *Macaca sylvanus* (Teufel et al, 2010),  [Diana’s monkey](https://pdf.sciencedirectassets.com/272524/1-s2.0-S0003347200X02308/1-s2.0-S0003347204002398/main.pdf?X-Amz-Security-Token=IQoJb3JpZ2luX2VjEBsaCXVzLWVhc3QtMSJIMEYCIQCplNmNzW58WwNzqyp7BJ28wFlx8TtjkjrSpJikDI5HVQIhAOV6tA6NPOqj%2BM%2FZNjSmO%2BMbjd5i26sCEhVO%2BQ6qCC8kKrwFCOT%2F%2F%2F%2F%2F%2F%2F%2F%2F%2FwEQBRoMMDU5MDAzNTQ2ODY1IgyHB4SJr09L0ahTuesqkAWNZ1qeXb6tiMwusKXHdZSYERQs8X%2FfFL72t7yx5qFhL%2FMzAc4JWYulgenzApF6tlulG1WzZ7E48awKg1Nahqz6XJ%2FyE1k3BVFed7ZvKRsGg%2Bp5XohzSNZHAwtk2mDTwdFDf5ON0k1HCMft9gU0Xaj4I%2B5gJgstmearfw5zPjfsR717fQfeMj0kyL0YtNp4NemzqQDiedl%2B4PjfqtL53yO04aV3bXmS6CwTpTlcLFKfoNKgjwkG1EqjD%2FsHvFQfUt%2FDM27hQNctQXxe1LkjHI5aTnjfbtJFuyXit5eSWdtsS%2FsZN41felLcRyNyHkKA2L5cQBMTRGY%2FV1tfmbTOq%2FFORRkt2VYwRksvfWvotkNDIQ2jt%2Fnx1aq9VqBKkcd%2B2rVfMtOv0FLD5ea6S9ZQqJXRf9t%2FIIalxWLrZQSAq%2F3BwCpMbkWtUqHppAZ%2FMRQ77PHRUSR2Wcow8HLfRcCxsrtL%2FVHcuLSTIDuZLZjU1TCXdPxrVjCXJ72ZPBOmfrOwYIQo%2FNg7QupNRQslSjrL2%2BI9GoCQS5h65CydRlyJA9uh8Fp4T53dNyOMdYx1YvTsSRI4hQVyHjmfK2on1wxdok%2FHRCSDHGrSR4dstviDJw5IaVcsRMjjDj38ze9Y%2Bw0BQzD%2FH%2BurPvRWwTle4%2B9i2dpyZ%2BQ0uvQjIbajpoTy8mzt4TPk50xd9sVZDa1blquW2jz8Rj3ZE7viAKcBsNNeVZZG6eUnodkqrhSCnXqSvyv58ESLHCVv0J7X6ws2Hpn9oz3%2BGMan%2F9z%2BkI6iRQVXPVqflAi%2FettSWave8%2BeAyyViCgTfqaezILkb8mpc%2BCefmQrY3OHQg3J0YTEs%2FMDIhpfhK4qQXJDSoltJ9nY73qHWYDDc2snLBjqwAUr7JoXyMRedQpBfFypiumr14LBHCL9RQ%2FJ9IHNv%2BEPy7UJhQSTaZZIoY5jQrCvYDVm689H76srn2%2Bynj2IR6YCB0sFx8uw6D8L9541spqRXgmDrQyMtEKosZIksDqY5QU0Ou2aKtc6LGQlHVnq5nLdrx9QSYGyitlZ3wc0rhQZFmxX%2BzGBQAucxO3%2Fh1yaYdrzJAcwSy9YMtO2o%2FTsD2mOcw%2FWioBL5IHtuKdUx6CoJ&X-Amz-Algorithm=AWS4-HMAC-SHA256&X-Amz-Date=20260122T192054Z&X-Amz-SignedHeaders=host&X-Amz-Expires=300&X-Amz-Credential=ASIAQ3PHCVTYUHV2GQ2S%2F20260122%2Fus-east-1%2Fs3%2Faws4_request&X-Amz-Signature=2091b9c5c14dd60f6c2b558675cf2e830687f5e3146ed3d65a2b217ed46eb018&hash=f3f55143595189c79d73ea4ebb5fe5e63a2ddaee35cb65fcaf5e6e07d4ee6c58&host=68042c943591013ac2b2430a89b270f6af2c76d8dfd086a07176afe7c76c2c61&pii=S0003347204002398&tid=spdf-45bd6f72-814f-41a0-9760-f3f4f8e66a7e&sid=d87edfd3113860410819056112234a39cac5gxrqb&type=client&tsoh=d3d3LnNjaWVuY2VkaXJlY3QuY29t&rh=d3d3LnNjaWVuY2VkaXJlY3QuY29t&ua=01065701505452585602&rr=9c21694d7e07654c&cc=gb), *Cercopithecus diana diana* (Scerif et al, 2004),  [Cotton top tamarin](https://psycnet.apa.org/fulltext/2002-00850-001.html), *Saguinus oedipus* (Neiworth et al, 2002),  [Olive baboon](https://onlinelibrary.wiley.com/doi/10.1002/ajp.22580), *Papio anubis* (Parron & Meguerditchian, 2016) | [Tufted capuchin](https://onlinelibrary.wiley.com/doi/abs/10.1002/ajp.23251?casa_token=Lx9iFCU8zzgAAAAA:_KNAUOUcRg6Dcny37THEr05efs54LvKxd3aodWQbCKdM2A8xJLpKpw9P-zONdaOsYovBtkAf5fz5CIo), *Sapajus libidinosus* (Falótico et al, 2021),  Bearded capuchin *Cebus libidinosus* (Mannu & Ottoni, 2009),  [Cottontop tamarin](https://www.sciencedirect.com/science/article/pii/S000334720293068X), *Saguinus oedipus* (Hauser et al, 2002),  [Long-tailed macaque](https://psycnet.apa.org/fulltext/2017-12252-001.html), *Macaca fascicularis* (Tan, 2017) |
| Other mammals | [Domestic dog](https://psycnet.apa.org/buy/1994-43961-001), *Canis familiaris* (Gagnon & Doré, 1994),  Domestic cat, *Felis catus* (Dumas & Doré, 1989) | [Domestic dog](https://www.science.org/doi/full/10.1126/science.1072702) (Hare et al, 2002),  Wolf, *Canis lupus* (Range & Virányi, 2011),  Silver fox, *Vulpes vulpes* (Hare et al, 2005),  [Dingo](https://link.springer.com/article/10.1007/s10071-009-0287-z), *Canis dingo* (Smith & Litchfield, 2009),  [Domestic goat,](https://www.sciencedirect.com/science/article/pii/S0003347204003264?via%3Dihub) *Capra hircus* (Kaminski et al, 2005),  [Domestic pig](https://brill.com/view/journals/beh/138/11-12/article-p1337_2.xml), *Sus scrofa domesticus* (Byrne et al, 2001) | Sea otter, *Enhydra lutis* (Payne & Jameson, 1984),  Bottlenose dolphin, *Tursiops sp*. (Mann et al, 2008) |
| Other |  | [Leopard gecko,](https://link.springer.com/article/10.1007/s10071-018-1230-y) *Eublepharis macularius* (Simpson & O’Hara, 2018) |  |
